# The effect of environmental information on evolution of cooperation in stochastic games

**DOI:** 10.1101/2022.10.18.512560

**Authors:** Maria Kleshnina, Christian Hilbe, Štěpán Šimsa, Krishnendu Chatterjee, Martin A. Nowak

**Affiliations:** Institute for Advanced Study in Toulouse, Toulouse, France; Max Planck Research Group Dynamics of Social Behavior, Max Planck Institute for Evolutionary Biology, Plön, Germany; IST Austria, Klosterneuburg, Austria; Faculty of Mathematics and Physics, Charles University, Prague, Czech Republic; Department of Mathematics, Harvard University, Cambridge, USA, Department of Organismic and Evolutionary Biology, Harvard University, Cambridge, USA

**Author notes:** These authors contributed equally to this work (Maria Kleshnina, Christian Hilbe).

## Abstract

Many human interactions feature the characteristics of social dilemmas where individual actions can have consequences for the group and the environment. The feedback between behavior and environment can be studied with the framework of stochastic games. In stochastic games, the state of the environment can change, depending on the choices made by group members. Past work suggests that such feedback can reinforce cooperative behaviors. In particular, cooperation can evolve in stochastic games even if it is infeasible in each separate repeated game. In stochastic games, participants have an interest in conditioning their strategies on the state of the environment. Yet in many applications, precise information about the state could be scarce. Here, we study how the availability of information (or lack thereof) shapes evolution of cooperation. Already for simple examples of two state games we find surprising effects. In some cases, cooperation is only possible if there is precise information about the state of the environment. In other cases, cooperation is only possible if there is no information about the state of the environment. We systematically analyze all stochastic games of a given complexity class, to determine when receiving information about the environment is better, neutral, or worse for evolution of cooperation.

---

Cooperation can be conceptualized as an individually costly behavior that creates a benefit to others^1^. Such cooperative behaviors have evolved in many species, from uni-cellular organisms to mammals^2^. Yet they are arguably most abundant and complex in humans, where they form the very basis of society, institutions and families^3,4^. Humans often support cooperation through direct reciprocity^5^. Here, people preferentially help those who have been helpful in the past^6^. Such forms of direct reciprocity naturally emerge when groups are stable, and when cooperation yields substantial returns^7^. In that case, individuals readily learn to engage in conditional cooperation, using strategies like Tit-for-tat^8–11^, Win-Stay Lose-Shift^12,13^, or multiplayer variants thereof^14–16^. When everyone adopts these strategies, groups can sustain cooperation despite any short-run incentives to free ride^17,18^.

To describe direct reciprocity formally, traditional models of cooperation consider individuals who face the same strategic interaction (game) over and over again. The most prominent model of this kind is the iterated prisoner’s dilemma^8^. In this game, two individuals (players) repeatedly decide whether to cooperate or defect. While the players’ decisions may change from one round to the next, the feasible payoffs remain constant. Models based on iterated games have become fundamental for our understanding of reciprocity. However, they presume that interactions take place in a constant social and natural environment. Individual actions in one round have no effect on the exact game being played in future. In contrast, in many applications, the environment is adaptive, such as when populations aim to control an epidemics^19–21^, manage natural resources^22–24^, or mitigate climate change^25–27^. Changing environments in turn often bring about a change in the exact game being played. Such applications are therefore best described with models in which there is a feedback between behavior and environment. In the context of direct reciprocity, such feedbacks can be incorporated with the framework of stochastic games^28–30^.

In stochastic games, individuals interact over multiple time periods. In each time period, the players’ environment is in one of several possible states. This state can change from one time period to the next, depending on the current state, the players’ actions, and on chance. Changes of the state affect the players’ available strategies and their feasible payoffs. In this way, stochastic games are better able to describe social dilemmas in which individual actions affect the nature of group’s future interactions.

Crucially, however, previous evolutionary models of stochastic games presume that individuals are perfectly aware of the current state^31–33^. This allows individuals to coordinate on appropriate responses once the state has changed. In contrast, in many applications, any knowledge about the state of the environment is at best incomplete. Such uncertainties can in turn have dramatic effects on human behavior^34–37^. In the following, we explore how state uncertainty shapes the evolution of cooperation in stochastic games. To this end, we compare two distinct scenarios. First, we consider the case when individuals are able to learn the state of their environment and condition their decisions on the current state. We will refer to this case as the *full-information setting*. In contrast, in the second case, individuals are aware that they are engaged in a stochastic game but they either ignore or are unable to obtain information about the current state, leading to them making decisions unconditioned on the environment. We refer to this case as the *no-information setting*. To compare these two settings we focus on the simplest possible case, where two players may experience two possible states. Already for this elementary setup, we obtain an extremely rich family of models that gives rise to many different possible dynamics. Already here, we observe that conditioning strategies on the state information can have drastic effects on how people cooperate.

To quantify the importance of state information, we introduce a measure to which we refer as the *value of information*. This value reflects by how much the cooperation rate in a population changes by gaining access to information about the present state. When this value is positive, access to information makes the population more cooperative. In that case, we speak of a *benefit of information*. In general, it is also possible to observe negative values, in which case we speak of a *benefit of ignorance*. With analytical methods for the important limit of weak selection^38–40^, and with numerical computations for arbitrary selection strengths, we compare the value of information across many stochastic games. We identify settings where receiving information is better, neutral, or worse for the evolution of cooperation. Most often, information is highly beneficial. However, there are also a few notable exceptions in which populations can achieve more cooperation when they are ignorant of their state. In the following, we describe and characterize these cases in detail.

## Results

### Stochastic games with and without state information

To explore the dynamics of cooperation in variable environments, we consider stochastic games^31–33^. We introduce our framework for the most simple setup, in which the game takes place among two players who interact for infinitely many rounds; some generalizations are discussed in the **SI**. In each round, players can find themselves in two possible states, *S* = {*s*_1_, *s*_2_}. Depending on the state, players engage in one of two possible prisoner’s dilemma games. In either game, they can either cooperate (*C*) or defect (*D*). Cooperation means to pay a cost *c* for the other player to get a benefit *b*_*i*_. The cost of cooperation is fixed, but the benefit *b*_*i*_ depends on the present state *s*_*i*_ (**Fig. 1a**). Without loss of generality, we assume that the first state is more profitable, such that *b*_1_ ≥ *b*_2_ *> c* := 1. However, states can change from one round to the next, depending on the game’s transition vector

**Fig. 1:**
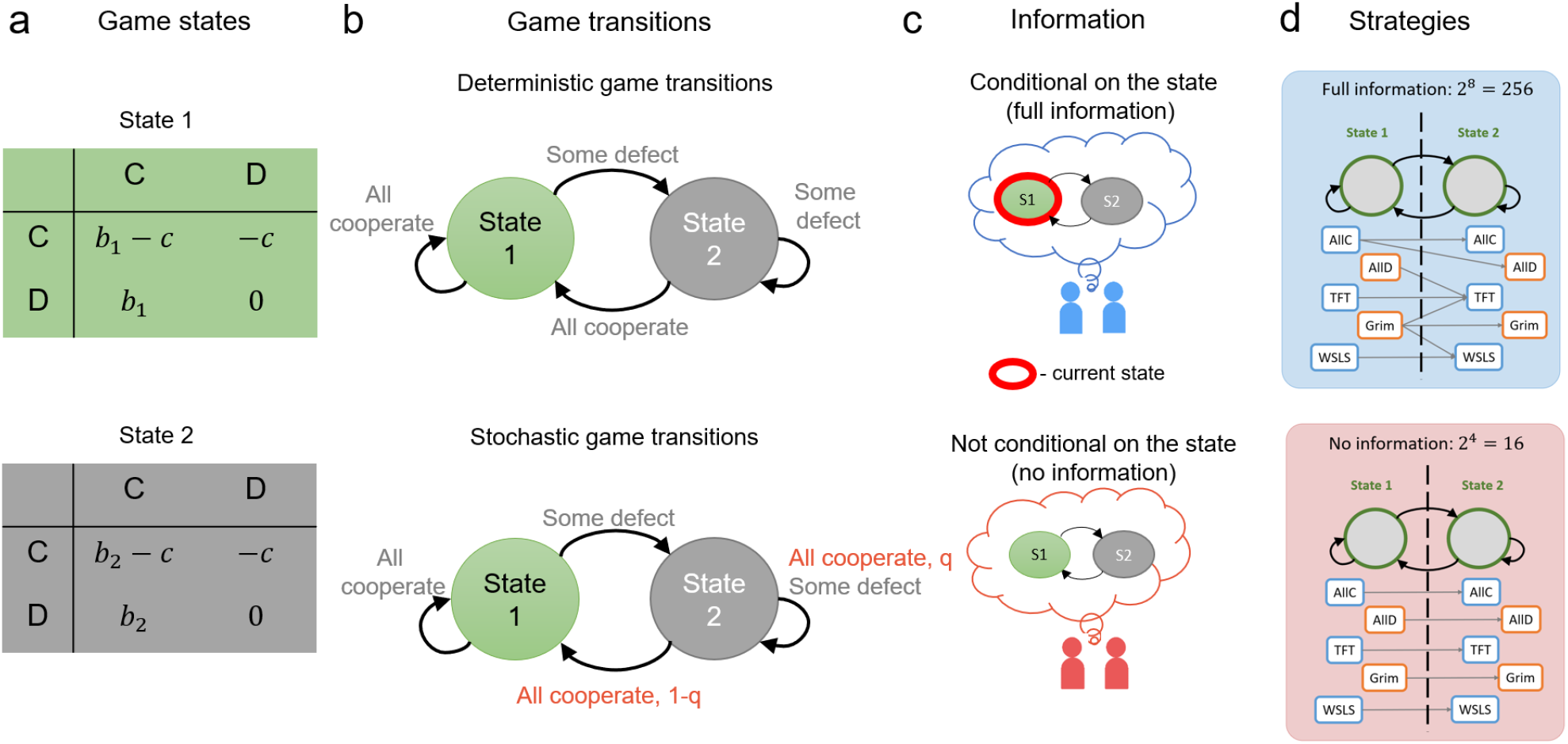
Stochastic games with full and no information. (a) We study 2-state stochastic games where transitions between the states depend on the actions of players. Each state is a Prisoners’ Dilemma with benefit *b*_1_ (or *b*_2_) and cost *c*. For instance, games in states 1 and 2 could correspond to some environmental conditions encoded in *b*_1_ and *b*_2_, respectively. We assume that *b*_1_ *> b*_2_. (b) Transitions between the states can be either completely determined by the actions of players in the current round (deterministic game transitions) or stochastic (almost-deterministic game transitions). In the latter, a game transition vector contains one stochastic transition *q*, which is the probability of the corresponding transition. For example, if some players defect in state 1 Prisoners’ Dilemma, then environmental conditions could worsen reducing the benefit. However, mutual cooperation might improve environmental conditions and recover a higher benefit. (c) Next, we consider two possible scenarios: when players condition their strategies on the information about the current state (for brevity, further referred to as “full info”) and when players do not condition their strategies on the information about the current state (for brevity, further referred to as “no info”). (d) In the case with full information, there are 2^8^ = 256 strategies, while without information there are only 2^4^ = 16 strategies. Players with full information can condition their strategies on the state of the environment and as a result play different strategies in different states. Players without information cannot condition their strategies and, hence, can play only one strategy in both environmental states.

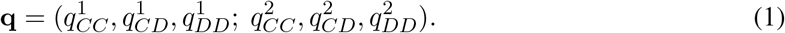

Here, each entry 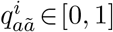 is the probability that players find themselves in the more profitable state *s*_1_ in the next round. This probability depends on the previous state *i* and on the players’ previous actions *a* and *ã*. For example, the transition vector **q** =(1, 0, 0; 1, 0, 0) corresponds to a game in which players are only in the more profitable state if they both cooperated in the previous round. Note that we assume the transition vector to be symmetric. That is, transition probabilities depend on the number of cooperators, but they are independent of who cooperated (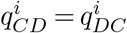 for all *i*). We say a transition vector is deterministic if each entry 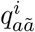 is either zero or one (**Fig. 1b**). Hence, there are 2^6^ = 64 deterministic transition vectors in total. We call a transition vector almost-deterministic if there is only one entry that is strictly between zero and one.

To explore how often players cooperate, depending on the information they have, we compare two settings (**Fig. 1c**). In the full-information setting, players learn the present state before making decisions. Thus, their strategies may depend on both the present state and on the players’ actions in the previous rounds. Herein, we assume that players make decisions based on memory-1 strategies, which only take into account the outcome of the last round^41^ (extensions to more complex strategies^42–46^ are possible, but for simplicity we do not explore them here). In the full information setting, memory-1 strategies take the form of an 8-tuple,

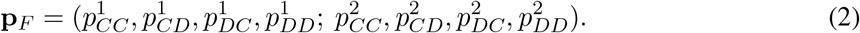

Here, 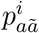 is the player’s probability to cooperate in state *i*, depending on the focal player’s and the co-player’s previous actions *a* and *ã*, respectively. We compare this full-information setting with a no-information setting, in which individuals are unable to condition their behavior on the current state. In that case, strategies are 4-tuples

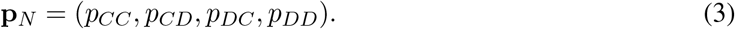

We note that the set of no-information strategies is a strict subset of the full-information strategies (they correspond to those **p**_*F*_ for which 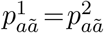 for all actions *a* and *ã*). Strategies (in either setting) are called deterministic if each entry is either zero or one. For full information, there are 2^8^ = 256 deterministic strategies. For no information, there are 2^4^ = 16 deterministic strategies.

The players’ strategies may be subject to errors with some small probability *ε*. In case of an error, an intended cooperation is misimplemented as a defection (and vice versa). As a consequence, a player with strategy **p** effectively uses strategy (1 − *ε*)**p** + *ε*(**1** − **p**). Given the error probability, the players’ strategies, and the game’s transition vector, we can compute how often players cooperate on average and which payoffs they get (see **Methods** for details).

Because we are interested in how cooperation evolves, we do not consider players with fixed strategies. Rather players can change their strategies in time, depending on the payoffs they yield. To describe this evolutionary dynamics, we use a pairwise comparison process^47^. This process considers populations of fixed size *N*. Players receive payoffs by interacting with all other population members. At regular time intervals, one player is randomly chosen and given the opportunity to revise its strategy. The player may do so in two ways. With probability *μ*, the player switches to a random deterministic memory-1 strategy (similar to a mutation in biological models of evolution). Otherwise, with probability 1−*μ*, the focal player compares its own payoff *π* to the payoff 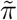 of a random role model. The player switches to the role model’s strategy with probability 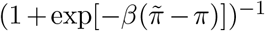. Here, *β >* 0 is the strength of selection. The higher the strength of selection, the more individuals are prone to imitate only those role models with a high payoff.

We study this evolutionary process analytically when mutations are rare and selection is weak (that is, when *μ, β* → 0). In addition, we numerically explore the process for arbitrary selection strengths. In either case, we compute which payoffs players receive on average and how likely they are to cooperate over time. By comparing the cooperation rates *γ*_*F*_ and *γ*_*N*_ for populations with full and no information, respectively, we quantify how favorable information is for the evolution of cooperation. We refer to the difference, *V*_*β*_(**q**) := *γ*_*F*_ − *γ*_*N*_ as the value of (state) information. In general, this value depends on the game’s transition vector **q**, as well as on the strength of selection *β*. When this value is positive, populations achieve more cooperation when they are able to learn the present state of the stochastic game. For more details, see **Methods**.

### The effect of state information in two examples

To begin with, we illustrate the effect of state information by exploring the dynamics of two examples. Both examples are variants of models that have been previously used to highlight the importance of stochastic games for the evolution of cooperation^31^. In the first example (**Fig. 2a**), players only remain in the more profitable first state if they both cooperate. If either of them defects, they transition to the inferior second state. Once there, they transition back to the more profitable state after one round, irrespective of the players’ actions. The second state may thus be interpreted as a ‘time-out’^31^. For numerical results, we assume that cooperation yields an intermediate benefit in the more profitable state and a low benefit in the inferior state (*b*_1_ = 1.8, *b*_2_ = 1.3).

**Fig. 2:**
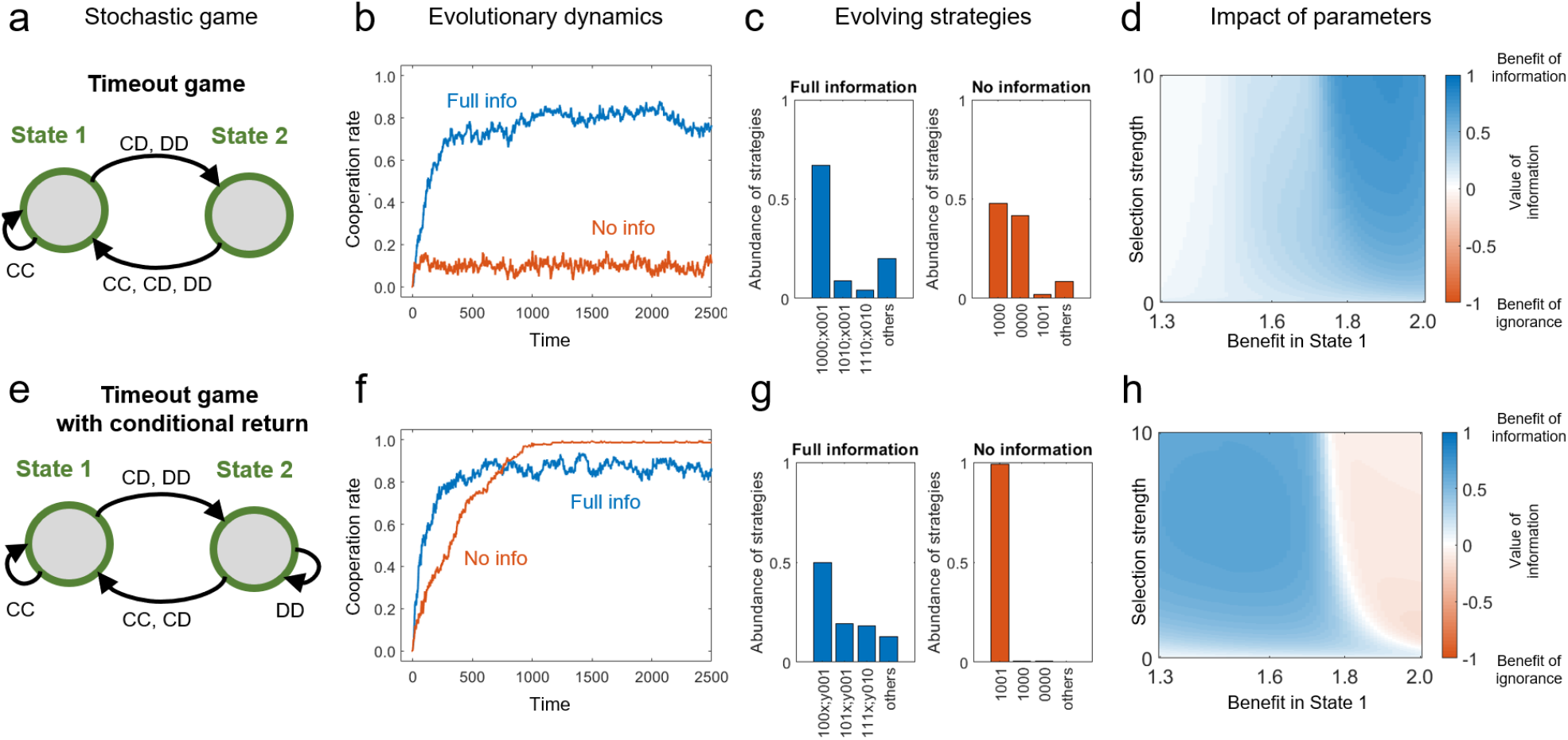
A comparison of the value of information in two games q_1_ = (1, 0, 0; 1, 1, 1) and q_2_ = (1, 0, 0; 1, 1, 0). (a-b) Game transitions for the timeout game q_1_ = (1, 0, 0; 1, 1, 1) and the timeout game with conditional returns **q**_2_ = (1, 0, 0; 1, 1, 0). (c-d) Evolutionary dynamics for the games with conditioning on the information about the environmental state (blue) and without conditioning (red). (e-f) Strategy abundance for games with full and no info for *β* = 10. Here, strategies are such that *x, y* ∈{ 0, 1}. (g-h) The impact of selection strength, *β*, and the benefit in state 1, *b*_1_, on the value of information. Parameter values: *b*_1_ = 1.8, *b*_2_ = 1.3, *c* = 1, population size *N* = 100, error rate *ε* = 0.01.

When we simulate the evolutionary dynamics of this stochastic game, we observe that individuals consistently learn to cooperate when they have full information. In contrast, without information, they mostly defect (**Fig. 2b**). To explain this result, we record which strategies are most likely to evolve in either case (**Fig. 2c**). In the full-information setting, individuals predominantly adopt a strategy **p**_*F*_ = (1, 0, 0, 0; *x*, 0, 0, 1), where *x* ∈ *{*0, 1*}* is arbitrary. This strategy may be considered as a variant of the win-stay lose-shift rule that has been successful in the traditional prisoner’s dilemma^12^. In particular, the strategy is fully cooperative with itself. Moreover, we prove in the **SI** that it is a subgame perfect (Nash) equilibrium if 2*b*_1_ − *b*_2_ ≥ 2*c* (this inequality is satisfied for the parameters we use) (**Fig. 3a**). On the other hand, in the no-information setting, this strategy is no longer available (**Fig. 3b**). Instead, players can only sustain cooperation with the traditional win-stay lose-shift (WSLS) rule **p**_*N*_ = (1, 0, 0, 1). This strategy is only an equilibrium under the more stringent condition *b*_1_ *>* 2*c*. Because our parameters do not satisfy this condition, cooperation does not evolve in the no-information setting. To explore how these results depend on the benefit of cooperation *b*_1_ and on selection strength *β*, **Fig. 2d** shows further simulations where we systematically vary both parameters. In all considered cases, state information is beneficial because it allows individuals to give more nuanced responses.

**Fig. 3:**
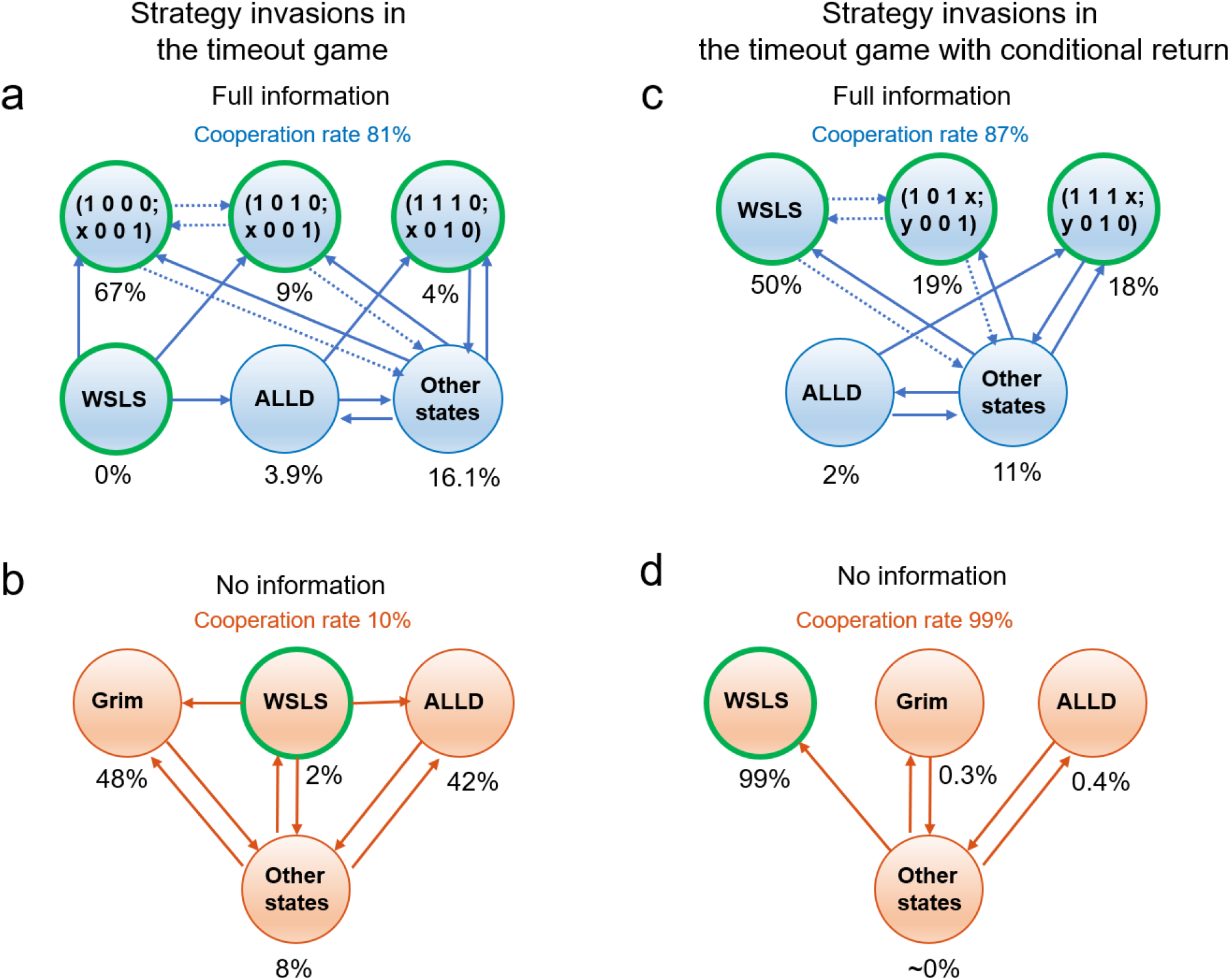
Strategy invasion analysis in two games q_1_ = (1, 0, 0; 1, 1, 1) >and q_2_ = (1, 0, 0; 1, 1, 0). We provide a strategy invasion analysis for two examples we considered in this manuscript: the game with a timeout in the state 2 (a-b) and the timeout game with conditional returns (c-d). Here, every circle represents a strategy evolving in this population with strategy frequency reported underneath the circle. Strategies that have 100% self-cooperation rate are highlighted with a green outline. Lines between the strategies represent the direction of selection: solid lines indicate that the fixation probability is larger than 1*/N*, whereas dotted lines indicate that the fixation probability is smaller than 1*/N* but greater than 1*/N ×* 10^−1^. Parameter values: *b*_1_ = 1.8, *b*_2_ = 1.3, *c* = 1, population size *N* = 100, error rate *ε* = 0.01.

The second example has a similar transition vector as the first, with a single modification. This time, the inferior state is only left if at least one of the two players cooperates (**Fig. 2e**). Although this modification may appear minor, the resulting dynamics is strikingly different. We observe that with and without state information, individuals are now largely cooperative. However, they are most cooperative when individuals do not condition their strategies on the state information (**Fig. 2f**). For this stochastic game, we show in the **SI** that already the traditional WSLS rule is a subgame perfect equilibrium for 2*b*_1_ −*b*_2_ ≥ 2*c*. As a result, WSLS is predominant in the no-information setting (**Fig. 3d**). In contrast, in the full-information setting, WSLS is subject to (almost) neutral drift by strategies that only differ from WSLS in a few bits (**Fig. 3c**). These other strategies may in turn give rise to the occasional invasion of defectors (**Fig. 2g**). Overall, we find that this stochastic game exhibits a benefit of ignorance when selection is sufficiently strong, and when cooperation is particularly valuable in the more profitable state (i.e., in the upper right corner of **Fig. 2h**).

These examples highlight three observations. First, just as there are instances in which state information is beneficial, there are also instances in which state information can reduce how much cooperation players achieve. Second, the stochastic games (transition vectors) for which state information is beneficial may only differ marginally from games with a benefit of ignorance. Finally, even if a stochastic game admits a benefit of ignorance, this benefit may not be present for all parameter values. Taken together, these observations suggest that in general, the effect of state information can be non-trivial and requires further investigation.

### A systematic analysis of the weak-selection limit

To explore more systematically in which cases there is a benefit of information (or ignorance), we study the class of all games with deterministic transition vectors. We first consider the limit of weak selection (*β* → 0). Here, game payoffs only weakly influence how individuals adopt new strategies. This limit plays an important role in evolutionary game theory more generally^38–40^. It often permits researchers to derive explicit solutions when analytical results are difficult to obtain otherwise. In our case, the limit of weak selection is particularly convenient, because it allows us to exploit certain symmetries between the two possible states (*s*_1_ and *s*_2_), and between the two possible actions (*C* and *D*, see **Methods** for details). As a result, we show that instead of 64 stochastic games, we only need to analyze 24. For each of these 24 transition vectors **q**, we explore whether information is beneficial, detrimental, or neutral (i.e., whether *V*_0_(**q**) is positive, negative, or zero).

First, we prove that half of the 64 stochastic games are neutral. In these games, the full-information and the no-information setting yield the same average cooperation rate. Among the neutral games, we identify three (overlapping) subclasses. (*i*) The first subclass consists of those games that have an absorbing state (15 cases). Here, either the first or the second state can no longer be left once it is reached (i.e., 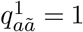 or 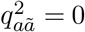, for all *a* and *ã*). For these games, state information is neutral because players can be sure they are in the absorbing state eventually. (*ii*) In the second subclass, transitions are state-independent^31^, which means 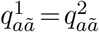 for all *a* and *ã* (6 additional cases). For deterministic transitions, state-independence implies that the current state can be directly inferred from the players’ previous actions, even without obtaining explicit state information. (*iii*) In the third subclass, neutrality arises because of more abstract symmetry arguments (see **Methods**). In particular, while the games in the previous two subclasses are neutral for all selection strengths, the games in the last subclass only become neutral in the limiting case of vanishing selection. One particular example of this last subclass is the game with transition vector **q** = (1, 0, 0; 1, 1, 0), which we studied in the previous section (**Fig. 2e–h**,**Fig. 3c–d**). There, we observed that this game can give rise to a benefit of ignorance when selection is intermediate or strong. Here, we conclude that this benefit disappears completely for vanishing selection (see also the lower boundary of **Fig. 2h**).

For the remaining 32 non-neutral cases, we identify a simple proxy variable that indicates whether or not the respective game exhibits a benefit of information for weak selection (**Fig. 4a**). Specifically, in a non-neutral game, information is beneficial if and only if *X >* 0, with *X* being

**Fig. 4:**
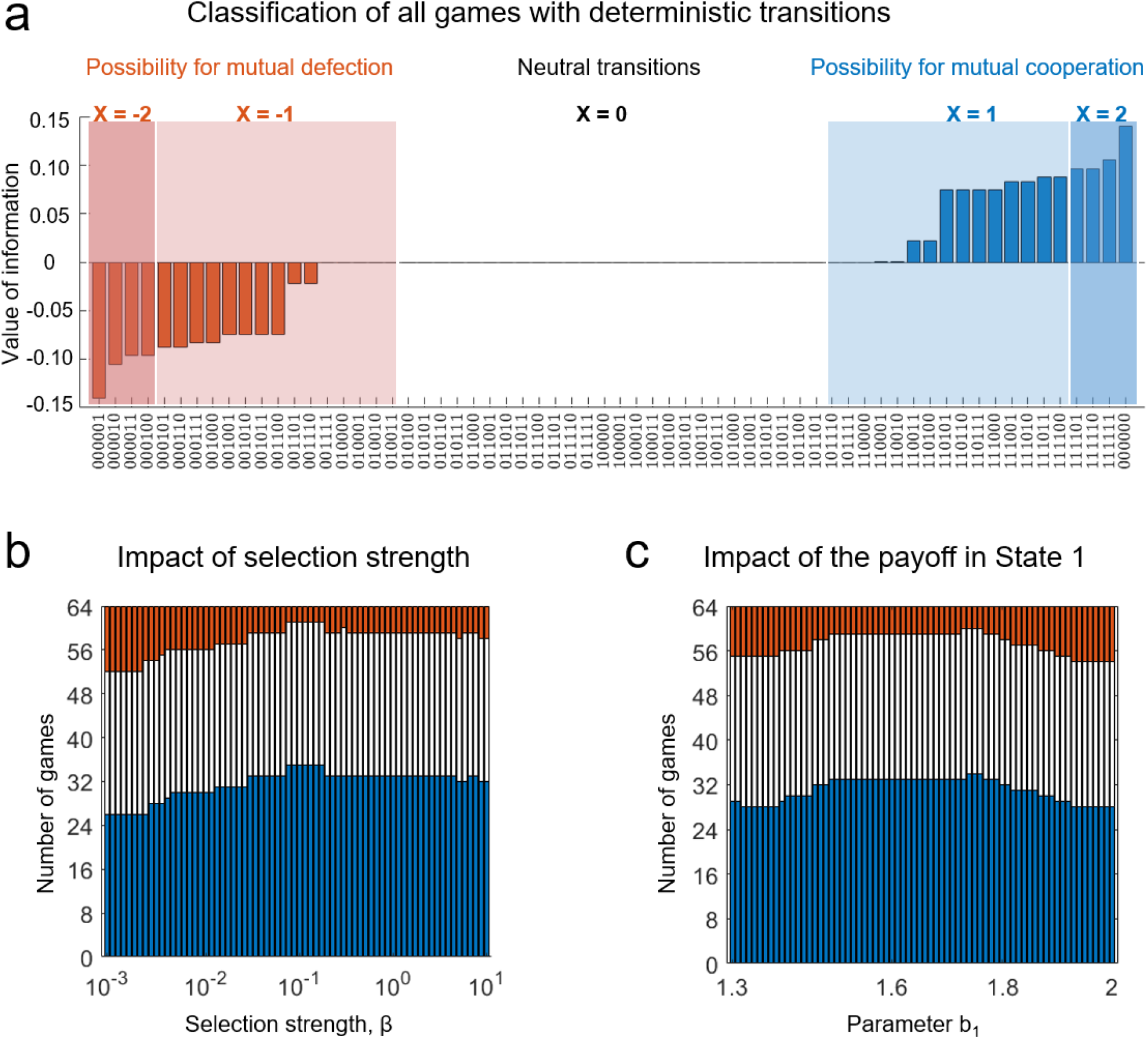
Classification of games with deterministic transitions. (a) We define a simple proxy variable, *X*, that indicates whether the stochastic game is absorbed in mutual cooperation or defection. A systematic analysis of all deterministic transition structures in the absence of selection. There are 64 transition vectors. For each of these vectors, we reproduce the value of information, *B*^*β*^(**q**), evaluated as a difference between cooperation rates for full and no information. No conditioning on the information always yields 50% cooperation. Conditioning on the information can yield the same cooperation rate (32 transition vectors), a lower cooperation rate (blue shaded, 16 vectors) or a higher cooperation rate (red shaded, 16 vectors). For games with a positive value of information, we obtain *X* ≥ 0. For games with a negative value of information, we obtain *X* ≤ 0. (b) This effect does not hold once selection strength increases. We calculate the value of information for different selection strengths, *β*. Every bar in the chart represents the value of *β* and colours represent the sign of the value of information across all 64 games. (c) We also study the effect of the benefit in State 1, *b*_1_, under the strong selection, *β* = 10. Parameter values: *b*_1_ = 1.8, *b*_2_ = 1.3, *c* = 1, population size *N* = 100, error rate *ε* = 0.01.

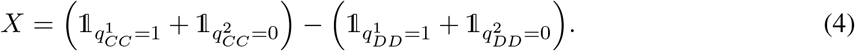

Here, 𝟙_*A*_ is an indicator function that is one if assertion *A* is true and zero otherwise. One can interpret the variable *X* as a measure for how easily the game can be absorbed in mutual cooperation (*X >* 0) or mutual defection (*X <* 0). For example, if a game has a transition vector with 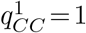, groups can easily implement indefinite cooperation by choosing strategies with 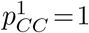. By doing so, players ensure they remain in the first state, in which they again would continue to cooperate. Using the proxy variable *X*, we can conclude that there are two properties of transition vectors that make state information beneficial in the limit of weak selection. The transition vector either needs to allow players to coordinate on mutual cooperation in a stable environment 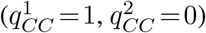; or it needs to prevent players from coordinating on mutual defection in a stable environment 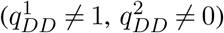. Again by symmetry considerations, we find that there are as many games with a benefit of information as there are games with a benefit of ignorance (16 cases each, see **Fig. 4b**).

### Exploring the impact of other game parameters

After characterizing the case of weak selection, we next explore the dynamics under strictly positive selection. To this end, we numerically compute the population’s average cooperation rate with and without state information, for each of the 64 stochastic games considered previously. To explore the impact of different game parameters, we systematically vary the strength of selection (**Fig. 4c, Fig. S1**), the benefit of cooperation (**Fig. 4b, Fig. S2**), and the error rate (**Fig. S3**). For 21 games, the evolving cooperation rates are the same with and without information. These games are neutral either because there is an absorbing state, or because transitions are state-independent (as described earlier). For the remaining cases, we find that a clear majority of them result in a benefit of information (**Fig. 4c,d**).

In the few cases in which we observe a consistent benefit of ignorance (the red squares in **Fig. S1–3**), there is overall very little cooperation. As a result, the magnitude of this benefit is negligible. Only in two cases one can find parameter combinations that lead to a sizeable benefit of ignorance for certain values of *b*_1_. The first such case is the stochastic game considered in **Fig. 2e–h** with transition vector **q** =(1, 0, 0; 1, 1, 0). The other case is only a slight modification of the first game, having a transition vector **q** =(1, 0, 1; 1, 1, 0). In both cases, players can sustain cooperation using the strategy WSLS even without state information, provided that 2*b*_1_−*b*_2_ ≥ 2*c*. But even when this condition holds, the benefit of ignorance is constrained, because even fully informed populations tend to achieve substantial cooperation rates (**Fig. S1–3**). Overall, these results suggest that among games with deterministic transitions, a sizeable benefit of ignorance is rare.

### The effect of environmental stochasticity

In our analysis so far, we assumed that the environment changes deterministically. Individuals who know the present state and the players’ actions can therefore anticipate the game’s next state. This form of predictability may overall diminish the impact of explicit state information because it reduces uncertainty. In the following, we extend our analysis to allow for stochasticity in the game’s transitions. To gain some intuition, we start with a simple example taken from the previous literature^31^ (see **Fig. 5a** for a depiction). According to the game’s transition vector, **q** = (1, 0, 0, *q*, 0, 0), players always find themselves in the less profitable second state if one or both players defect. If both players cooperate, however, they either remain in the first state (if they are already there), or they transition to the first state with probability *q* (if they start out in the second state). This stochastic game represents a scenario in which an environment deteriorates immediately once players defect. If players resume to cooperate, it may take several rounds for the environment to recover.

**Fig. 5:**
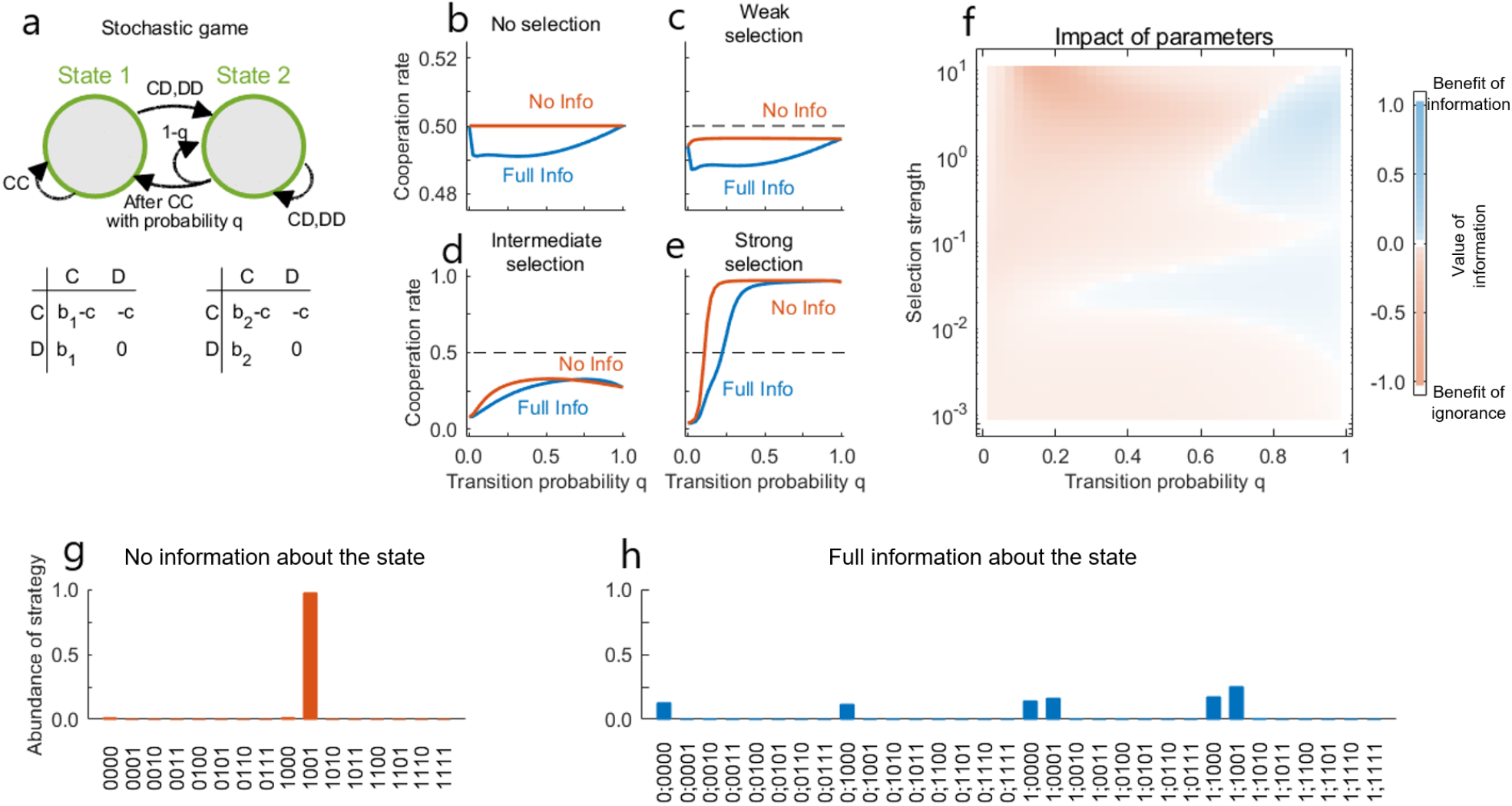
Benefit of ignorance in a game with one stochastic transition. (a) We consider a stochastic game in which defection by any player always leads to the second state. From there, they return to the first state after mutual cooperation with probability *q*. (b–f) We have computed numerically exact cooperation rates for the stochastic game without conditioning on the information (red) and with conditioning on the information (blue) for different values of the transition probability *q* and selection strength *β*. For no and weak selection, there is a benefit of ignorance for all values of *q* ∈ (0, 1) (that is, the no information scenario always yields more cooperation). But even for intermediate and strong selection, some benefit of ignorance may persist. (g,h) For *q* = 0.2 we observe in the limit of strong selection that most players adopt WSLS when players do not condition their strategies, whereas there is no clearly winning strategy in the case when players condition their strategies on the information about the environmental state. Parameter values: *b*_1_ = 1.8, *b*_2_ = 1.3, *c* = 1, error rate *ε* = 0.01. For no, weak, intermediate and strong selection we use *β* = 0, *β* = 0.001, *β* = 1, and *β* = 10, respectively.

For this example, we find that the value of information varies non-trivially, depending on the transition probability *q* and the strength of selection *β* (**Fig. 5b–e**). Overall, parameter regions with a benefit of ignorance seem to prevail (**Fig. 5f**). To obtain analytical results, again we study the game for weak selection (*β* → 0). In that case, the value of information can be computed explicitly, as 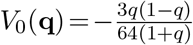. In particular, there is a benefit of ignorance for all intermediate values *q* ∈ (0, 1). This benefit becomes most pronounced for 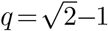 (for all formal derivations, see **SI**). As we increase the selection strength, however, the dynamics can change, depending on *q* (the probability to transition from the second state to the first after mutual cooperation). If *q* is large, there can be a (small) benefit of information. However, as *q* becomes smaller, there is a considerable benefit of ignorance (**Fig. 5f**).

To explore this benefit of ignorance in detail, we record which strategies players adopt for *q* = 0.2. Without state information, we find that players adopt WSLS almost all of the time (**Fig. 5g**). In contrast, when players condition their strategies in the state information, WSLS is risk-dominated by a strategy that has been termed Ambitious WSLS^31^ (AWSLS). AWSLS differs from WSLS after mutual cooperation, in which case AWSLS only cooperates when players are in the first state (i.e., 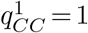 but 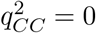). Once AWSLS is common in the population, it opens up opportunities for less cooperative strategies to invade. In particular, also non-cooperative strategies (like ALLD) are adopted for a non-negligible fraction of time (**Fig. 5h**, and **SI** for details). Overall, again we observe that predicting the effect of information is non-trivial. While some parameter combinations favor populations with full information, we also observe a benefit of ignorance for a significant portion of the parameter space.

To obtain a more comprehensive picture, we numerically analyze all stochastic games with almost-deterministic transition vectors. Because the corresponding transition vectors have exactly one entry *q* between 0 and 1, there are 6·2^5^ = 192 cases in total. We find several regularities. First, similarly to games with deterministic transitions, we find that there are 24 transition vectors for which the game is neutral. In all of the respective games, one of the two states is absorbing. Hence conditional and unconditional strategies perform in similarly. Second, we can analyze the remaining cases in the limit of vanishing selection (**Fig. S4**). Most of these games follow the rule defined by the proxy variable *X* in Eq. (4), with some exceptions discussed in detail in the **SI**. Finally, for all selection strengths we compute the players’ average cooperation rates numerically. We do this for all 192 games for weak (**Fig. S5**), intermediate (**Fig. S6**), and strong selection (**Fig. S7**). Similar to the case of deterministic transitions, state information is beneficial in an absolute majority of cases (**Fig. S8**). However, exceptions can and do occur. A benefit of ignorance arises most frequently when mutual cooperation in a more beneficial state results in the game remaining in a more profitable state, and when mutual defection in any state is punished with deteriorating environmental conditions.

The method we propose here is not limited to only specific transition vectors we demonstrated so far. To obtain a better understanding of the general effect of the information about the state of the environment, we numerically study a wider range of transition vectors. We assume the entries of **q** come from a finite grid 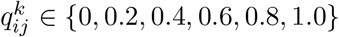. In this example, there are 6^6^ = 46, 656 stochastic games, the majority of which fall into the category of games where the state information is beneficial for cooperation (**Fig. S9b**). Yet, there is a non-negligible number of games where strategies unconditioned on the environmental state result in higher cooperation rates. However, generally the benefit of ignorance is smaller than the benefit of information (**Fig. S9a**).

## Discussion

When people interact in a social dilemma, their actions often have spillovers to their social, natural, and economic environment^48–51^. Changes in the environment may in turn modulate the characteristics of the social dilemma. One important example of such a feedback loop is the ‘tragedy of the commons’^52^. Here, groups with little cooperation may deteriorate their environment, thereby restricting their own feasible long-run payoffs.

Such spillovers between the groups’ behavior and their environment can be formalized as a stochastic game^28^. In stochastic games, individuals interact for many time periods. In each time period, they may face a different kind of social dilemma (state). The way they act in one state may affect the state they experience next. Recently, stochastic games have become a powerful model for the evolution of cooperation, because changing environments can reinforce reciprocity^31–33^. In particular, the evolution of cooperation may be favored in stochastic games even if cooperation is disfavored in each individual state^31^. However, implicit in these studies is the assumption that individuals are perfectly aware of the state they are in. Here, we systematically explore the implications of this assumption. We study to which extent individuals learn to cooperate, depending on whether or not they condition their strategies on the state information in a stochastic game. Alternatively, this setup can be interpreted as a game where players do not know that they are engaged in a stochastic game. In such case, players learn their payoffs only at the end of the game and, hence, are unaware of the environmental transitions. In this manuscript, we mostly focused on the first interpretation. We say the stochastic game shows a benefit of information if well-informed groups tend to be more cooperative. Otherwise, we speak of a benefit of ignorance.

Already for the most basic instantiation of a stochastic game, with two individuals and two states, we find that the impact of information is non-trivial. All three cases are possible: state information can be beneficial, neutral, or detrimental for cooperation. To explore this complex dynamics, we employ a mixture of analytical techniques and numerical approaches. Analytical results are feasible in the important limiting case of weak selection^38–40^. Here, we observe an interesting symmetry. For every stochastic game in which there is a benefit of information, there is a corresponding game with a benefit of ignorance. This symmetry breaks down for positive selection. As selection increases, we observe more and more cases in which state information becomes beneficial. Moreover, in those few cases in which a benefit of ignorance persists, this benefit tends to be small. These results highlight the importance of accurate state information for responsible decision making.

However, there are a few notable exceptions. In particular, we identify several ecologically plausible scenarios where individuals cooperate more when they ignore their environment’s state. One example is the game displayed in **Fig. 2e–h**. Here, players only remain in the profitable state when they both cooperate. Once they defect, they transition to the less profitable state. From there, they can only escape if at least one player cooperates. This game reflects a scenario where the group’s environment further reinforces cooperation. Cooperative groups are rewarded by maintaining access to the more profitable state. Non-cooperative groups are punished by transitioning to an inferior state. For this kind of environmental feedback it was previously observed that the simple win-stay lose-shift (WSLS) strategy can sustain cooperation easily^31–33^. WSLS can be instantiated without any state information. Once a population settles at WSLS, providing state information can be harmful: it permits individuals to deviate towards more nuanced strategies, and hence it undermines overall cooperation.

To allow for a systematic treatment, we focus here on comparably simple games. Nevertheless, the number of games we consider is huge. For example, if all transitions between states are assumed to be deterministic (independent of chance), there are 64 cases to consider (**Figures S1–S3**). If all but one transition are assumed to be deterministic, we obtain 192 families of games (each having a free parameter *q* ∈ [0, 1], **Figures S4–S8**). In all these instances, we observe that seemingly innocent changes in the environmental feedback or in the game parameters can lead to complex changes in the dynamics. In particular, games with a benefit of information may turn into games with a benefit of ignorance. These observations suggest that predicting the effect of state information is generally non-trivial. These difficulties are likely to increase as we extend the model to larger groups^53,54^, more complex strategies^42–46^, or environments with multiple possible states^31^.

While we systematically study the effect of state information in a simple two-player two-state setup, our framework is in fact more general. For example, **Fig. S10**illustrates the results of a public goods game among three players. Again, there are transitions vectors for which information proves to be beneficial. Yet, with a slight modification of the transition vector, we can observe a benefit of ignorance.

Here, we introduce a simple and easily generalizable framework to explore how state information (or the lack thereof) affects the evolution of cooperation. Depending on the exact environmental feedback, the effect can be beneficial, neutral, or detrimental. As a general rule of thumb, we observe that state information is highly valuable. Information is particularly relevant when selection is strong (such that payoffs have a notable impact on decisions). There are, however, a few notable exceptions, in which it is inconsequential or even advantageous if the group ignores the information about their state. Overall, these results further illustrate the intricate dynamics that arise in the presence of environmental, informational, and behavioral feedbacks. By exploring these feedbacks in elementary stochastic games, we can better understand the more complex dynamics of the socio-ecological systems around us.

## Methods

### Calculation of payoffs in stochastic games

In the following, we describe how we can calculate the payoffs when players with arbitrary full-information strategies interact with each other. Since any no-information strategy can be associated with a full-information strategy, the same method also allows us to compute payoffs when a full-information player is matched with a no-information player, or when two no-information players are matched.

We consider games that are infinitely repeated and in which there is no discounting of the future. Given the players’ effective memory-1 strategies **p** and 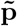, such games can be described as a Markov chain. The states of this Markov chain correspond to the eight possible outcomes *ω* = (*s*_*i*_, *a, ã*) of a given round. Here, *s*_*i*_ ∈ *{s*_1_, *s*_2_*}* reflects the environmental state, whereas *a, ã* ∈ *{C, D}* are player 1’s and player 2’s actions, respectively. The transition probability to move from state *ω* = (*s*_*i*_, *a, ã*) in one round to 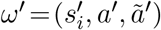 in the next round is given by a product of three factors,

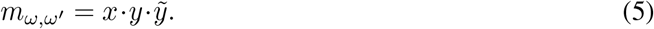

The first factor

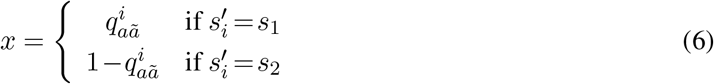

reflects the probability to move from environmental state *s*_*i*_ to 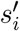, given the player’s previous actions. Since the game is symmetric, we note that 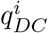 is defined to be equal to 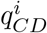. The other two factors are given by

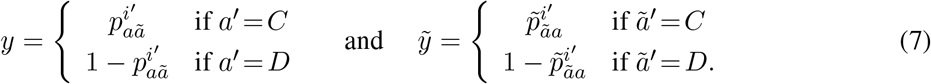

They correspond to the conditional probability that each of the two players chooses the action prescribed in *ω*^*t*^.

By collecting all these transition probabilities, we obtain an 8*×*8 transition matrix *M* =(*m*_*ω,ω*′_). Assuming that players are subject to errors and that the game’s transition vector satisfies **q ≠** (1, 1, 1, 0, 0, 0), this transition matrix has a unique left eigenvector **v**. The entries 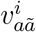 of this eigenvector give the frequency with which players observe the outcome *ω* = (*s*_*i*_, *a, ã*) over the course of the game. For a given transition vector **q**, we can thus compute the first players’ expected payoff as

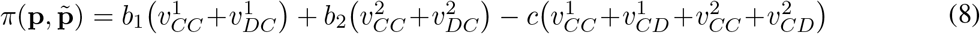

The second player’s payoff can be computed analogously. We define the average cooperation rate of the two players as

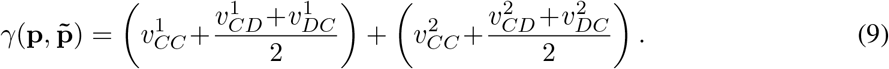

### Evolutionary dynamics

To model how players learn to adopt new strategies over time, we study a pairwise comparison process^47^ in the limit of rare mutations^55–58^.

We consider a population of fixed size *N*. Initially, all players adopt the same resident strategy **p**_*R*_ =*ALLD*. Then one of the players switches to a randomly chosen alternative strategy **p**_*M*_. This mutant strategy may then either go extinct or reach fixation eventually, depending on which payoff it yields compared to the resident strategy. If the number of players adopting the mutant strategy is given by *k*, the expected payoffs of the two strategies is

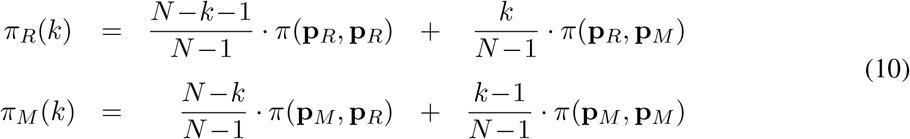

Based on the players’ payoffs, the fixation probability of the mutant strategy can be computed explicitly^38,59^. It is given by

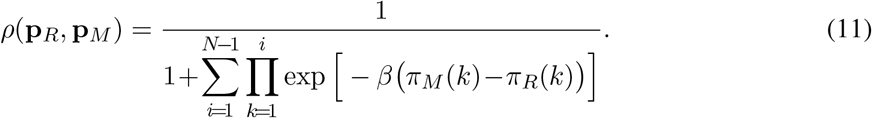

Here, *β* ≥ 0 is the strength of selection. It reflects how important relative payoff advantages are for the evolutionary success of a strategy. If there is no selection and *β* = 0, payoffs are completely irrelevant. In that case the fixation probability of a single mutant player in a resident population of size *N* simplifies to *ρ*(**p**_*R*_, **p**_*M*_) = 1*/N*. As *β* increases, the fixation probability is increasingly biased in favour of mutant strategies with a high relative payoff.

If the mutant fixes, it becomes the new resident strategy. Then another mutant strategy is introduced and either fixes or goes extinct. By iterating this basic process for *τ* time steps, we obtain a sequence (**p**_0_, **p**_1_, **p**_2_, …, **p**_*τ*_) where **p**_*t*_ is the resident strategy present in the population after *t* mutant strategies have been introduced. Based on this sequence, we can calculate the population’s average cooperation rate and payoff as

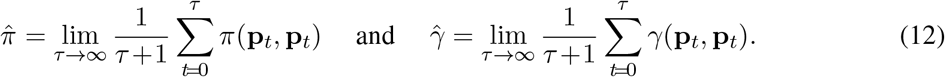

In general, these payoff and cooperation averages can only be approximated, by simulating the above described process for a sufficiently long time *τ*.

Numerically exact results are feasible when mutant strategies are taken from a finite set 𝒫. In that case, the evolutionary dynamics can again be described as a Markov chain^55^. Each state of this Markov chain corresponds to one possible resident population **p** ∈ *𝒫*. Given that the current resident population uses **p**, the probability that the next resident population uses strategy 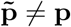 is given by 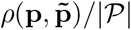. By calculating the invariant distribution **w** = (*w*_**p**_) of this Markov chain, we can compute the average cooperation rates and payoffs according to Eq. (12) by evaluating

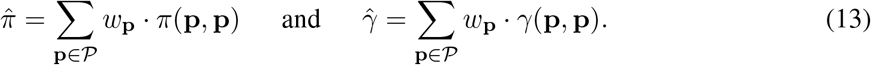

In this manuscript, we report results for the two specific strategy sets 𝒫_*F*_ and 𝒫_*N*_. By comparing the respective averages 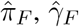 with the averages 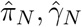, we wish to characterise for which stochastic games there is an advantage of not knowing the current environmental state.

### Transition vectors’ transformations

For each stochastic game **q**, we can define an associated <u>twin</u> *χ*(**q**) by relabelling the states,

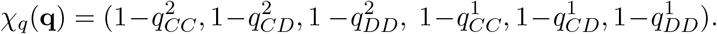

Second, for *β* = 0 it makes no difference to which behaviour we refer to as “Cooperation” or “Defection”. It follows that for each stochastic game **q** we can define an associated <u>mirror</u> game *ψ*(**q**) by flipping the meaning of C and D,

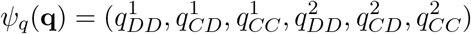

Of course, we can also consecutively perform both transformations. This yields a third transformation that we call a mirror-twin,

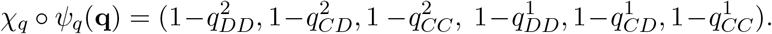

We define similar transformations *χ*_*p*_(**p**) and *ψ*_*p*_(**p**) for the players’ memory-one strategies, and transformations *χ*_*v*_(**v**) and *ψ*_*v*_(**v**) for the resulting stationary distributions.

If **v**(**p** | **q**) denotes the stationary distribution of the stochastic game **q** between two **p**-players, we show the following relations

i. *χ*_*v*_ **v**(*χ*_*p*_(**p**) *χ*_*q*_(**q**)) = **v p q** and *ψ*_*v*_ **v**(*ψ*_*p*_(**p**) *ψ*_*q*_(**q**)) = **v p q**
ii. *γ* **p q** = *γ χ*_*p*_(**p**) *χ*_*q*_(**q**) and *γ* **p q** = 1−*γ ψ*_*p*_(**p**) *ψ*_*q*_(**q**);
iii. Let 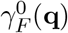 be the average cooperation rate in a full information game with transitions **q** and selection strength *β* = 0. Then

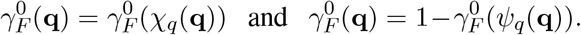

In general, we find that among the 64 deterministic games, exactly half of them is neutral (Fig. 3).

All these cases fall within four possible categories:

1. The transition vector has an absorbing state: 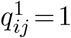 or 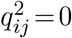 for all *i, j* ∈ *{C, D}*.
2. The transition vector is its own mirror: *ψ*_*q*_(**q**) = **q**.
3. The transition vector is its own mirror-twin: *χ*_*q*_ *? ψ*_*q*_(**q**) = **q**.
4. The transition vector is state-independent: 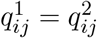 for all *i, j* ∈ *{C, D}*.

---

**Fig. S1:**
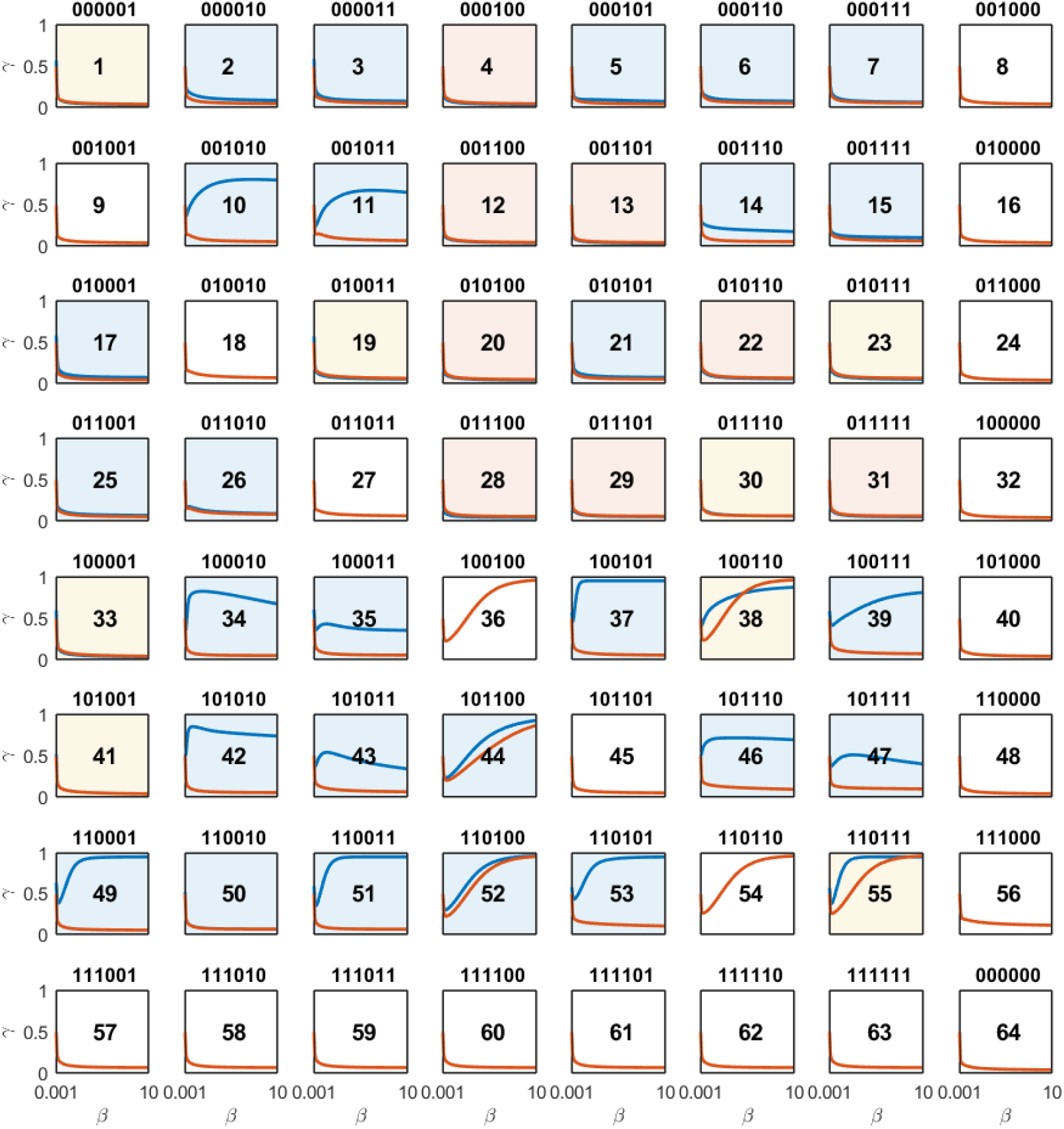
A systematic analysis of all deterministic transition structures for different selection strengths. We plot cooperation rates for the games with conditioning (blue line) and no conditioning (red line) on the information as functions of selection strength *β*. There are 2^6^ = 64 transition structures in which all transitions are either 0 or 1. For each of these transition structures, we reproduce the cooperation rates for both cases. Conditioning on the information can yield the same cooperation rate (white background, 21 instances), a higher cooperation rate (blue shaded, 27 instances), a lower cooperation rate (red shaded, 8 instances) or a change in the cooperation rates difference (yellow shaded, 8 instances). Games without any effect of information (white) have the same cooperation rates for any selection strength. Parameter values: *b*_1_ = 1.8, *b*_2_ = 1.3, *c* = 1, population size *N* = 100, error rate *ε* = 0.01.

**Fig. S2:**
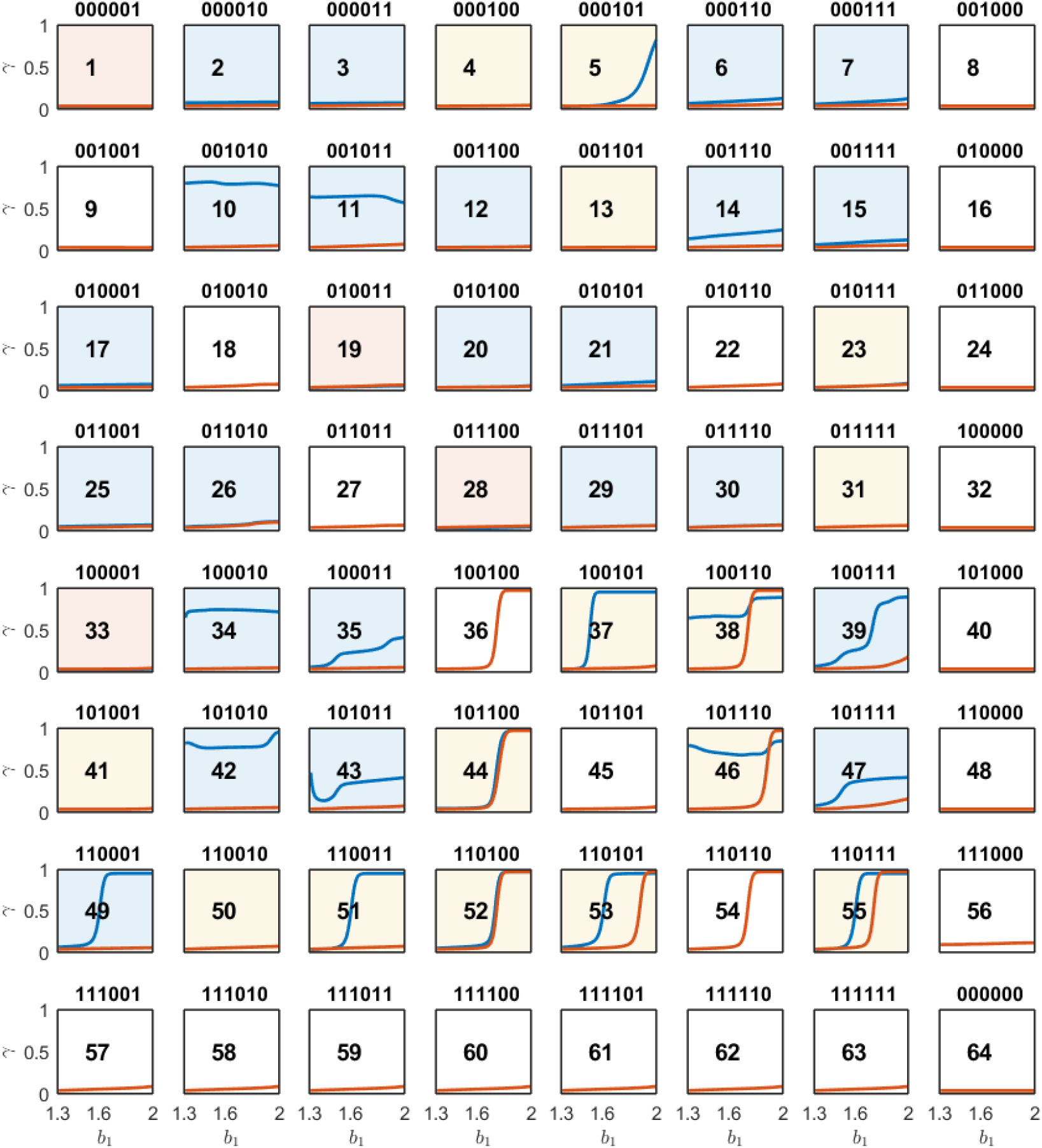
A systematic analysis of all deterministic transition structures for different benefit *b*_1_ and *β* = 10. Conditioning on the information can yield the same cooperation rate (white background, 22 instances), a higher cooperation rate (blue shaded, 23 instances), a lower cooperation rate (red shaded, 4 instances) or a change in the cooperation rates difference (yellow shaded, 15 instances). Games without any effect of information (white) have the same cooperation rates for any benefit *b*_1_. Parameter values: *b*_2_ = 1.3, *c* = 1, population size *N* = 100, error rate *ε* = 0.01.

**Fig. S3:**
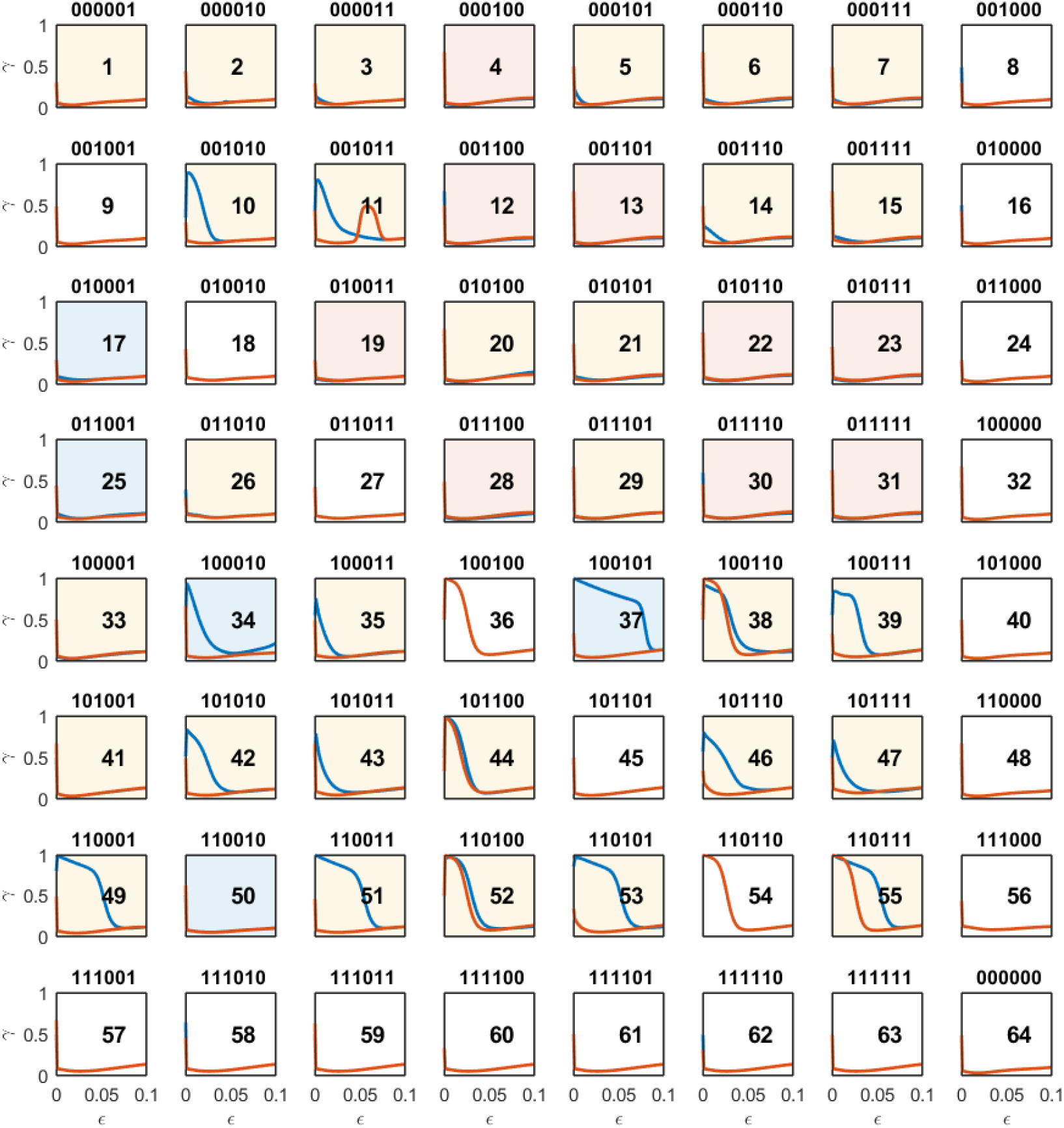
A systematic analysis of all deterministic transition structures for different error rate *ϵ* and *β* = 10. Conditioning on the information can yield the same cooperation rate (white background, 21 instances), a higher cooperation rate (blue shaded, 5 instances), a lower cooperation rate (red shaded, 9 instances) or a change in the cooperation rates difference (yellow shaded, 29 instances). Games without any effect of information (white) have the same cooperation rates for any cost of cooperation *ϵ*. Parameter values: *b*_1_ = 1.8, *b*_2_ = 1.3, *c* = 1, population size *N* = 100,.

**Fig. S4:**
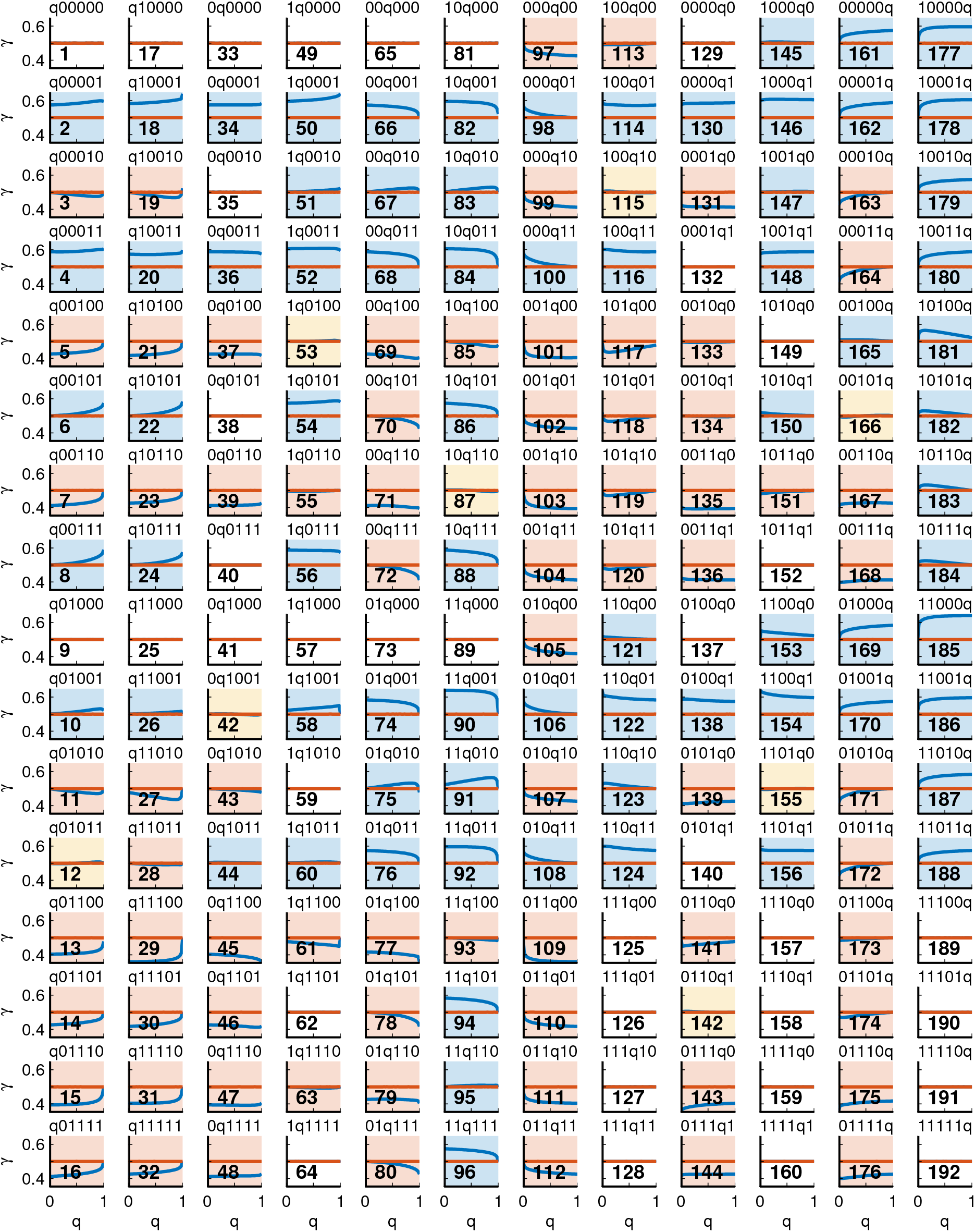
A systematic analysis of the almost-deterministic transition structures in the absence of selection. There are 6 2^5^ = 192 transition structures in which exactly one transition (one entry of **q**) is probabilistic. For each of these transition structures, we reproduce the cooperation rates for cases with and without conditioning on the environmental information. No conditioning always yields 50% cooperation. Conditioning can yield the same cooperation rate (white, 36 instances), a higher cooperation rate (blue, 74 instances) or a lower cooperation rate (red, 74 instances). Parameter values: *b*_1_ = 1.8, *b*_2_ = 1.3, *c* = 1, population size *N* = 100, error rate *ε* = 0.01.

**Fig. S5:**
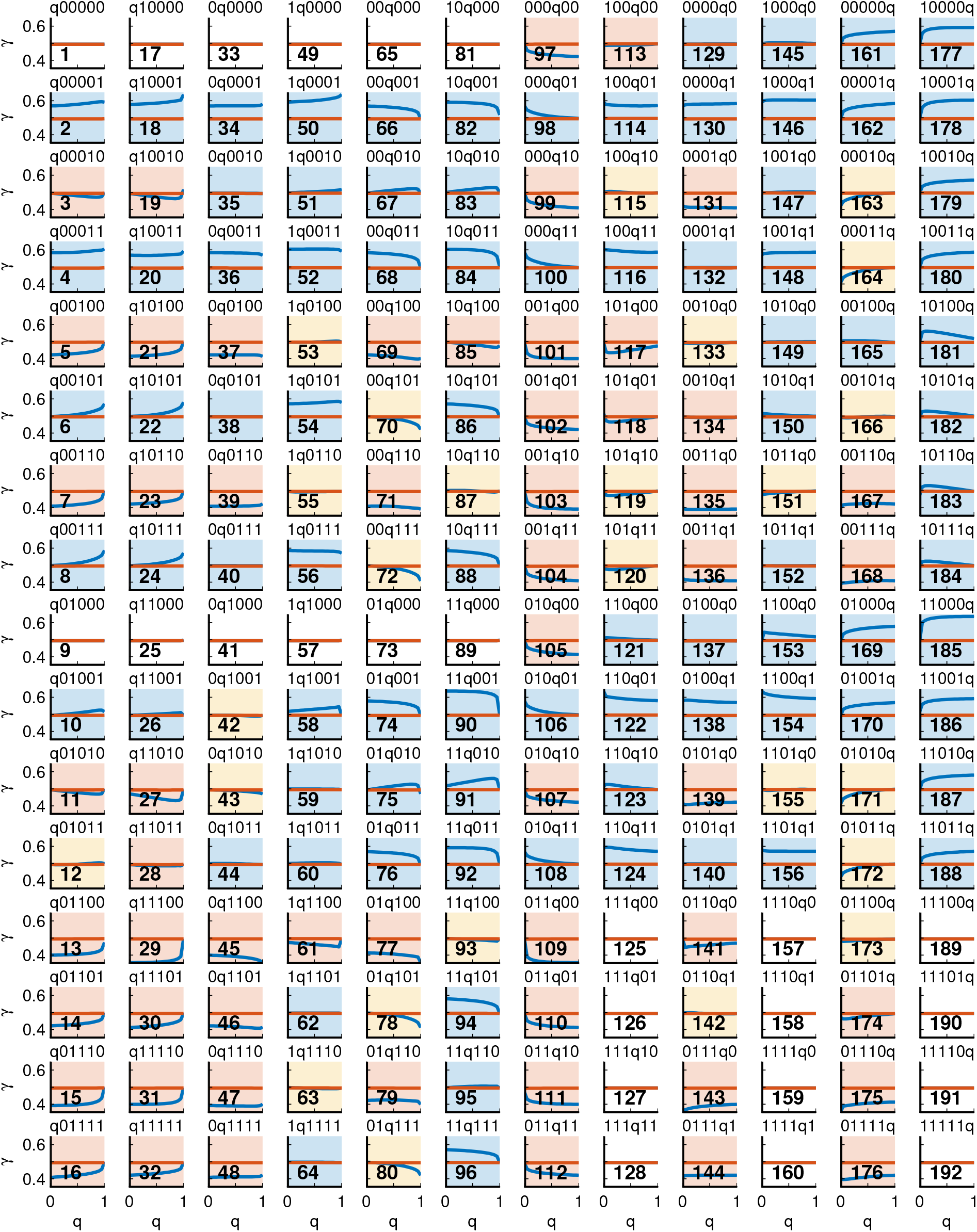
Almost-deterministic transition structures for weak selection. Same as Figure 2, but for *β* = 0.001 instead of *β* = 0. Parameter values: *b*_1_ = 1.8, *b*_2_ = 1.3, *c* = 1, population size *N* = 100, error rate *ε* = 0.01.

**Fig. S6:**
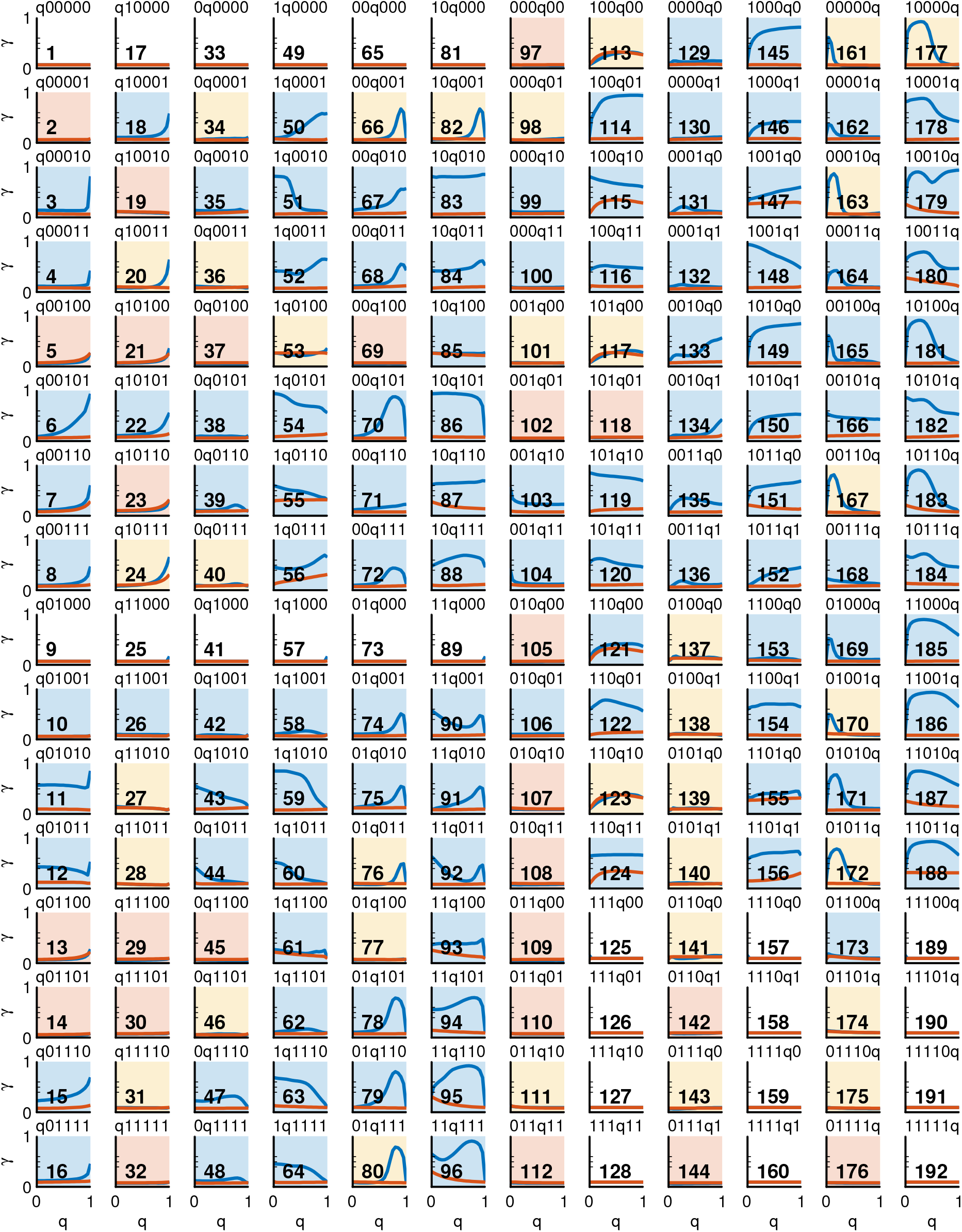
Almost-deterministic transition structures for intermediate selection. Same as previous two figures, but using *β* = 1. Parameter values: *b*_1_ = 1.8, *b*_2_ = 1.3, *c* = 1, population size *N* = 100, error rate *ε* = 0.01.

**Fig. S7:**
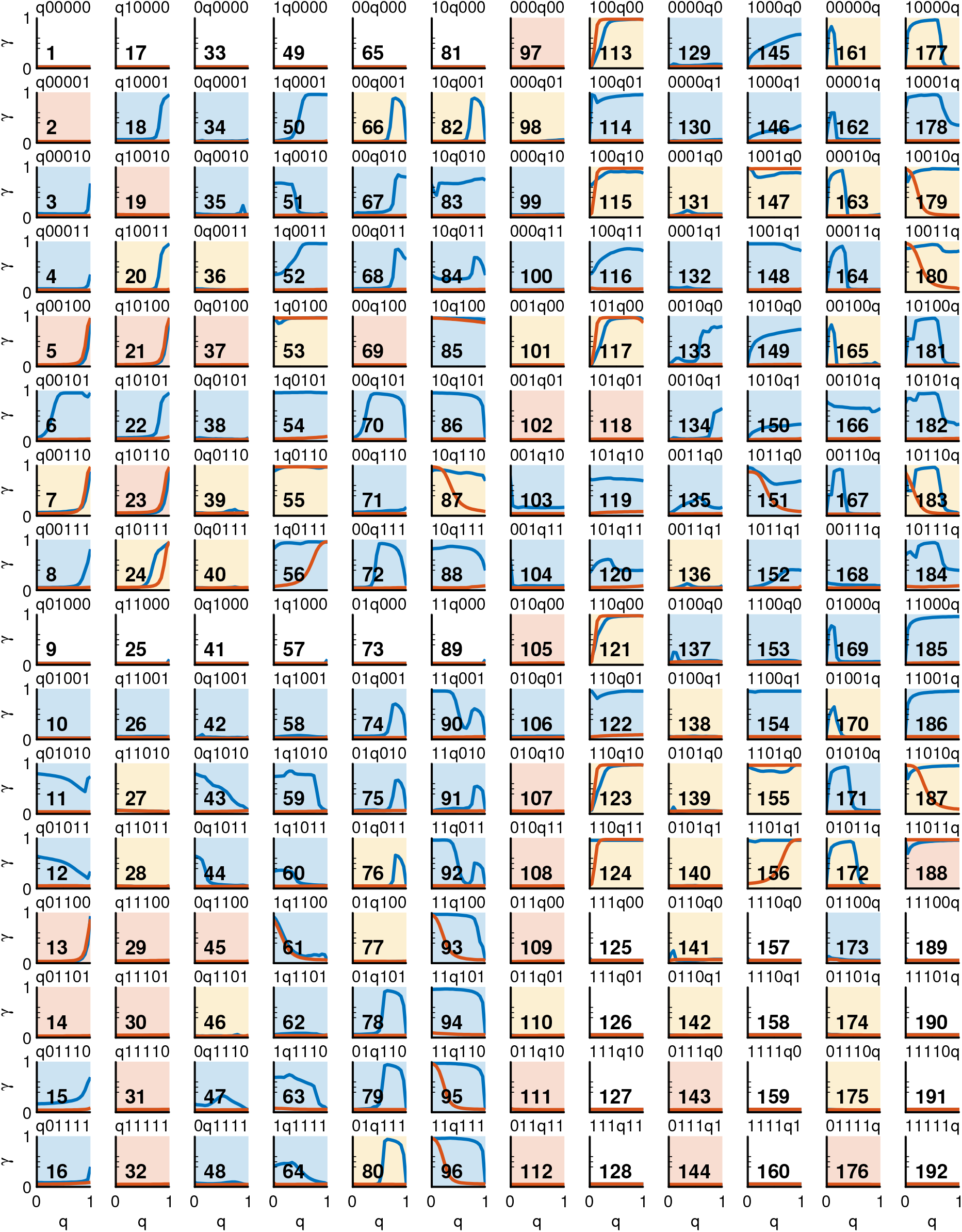
Almost-deterministic transition structures for strong selection. Same as previous figures, but using *β* = 10. Parameter values: *b*_1_ = 1.8, *b*_2_ = 1.3, *c* = 1, population size *N* = 100, error rate *ε* = 0.01.

**Fig. S8:**
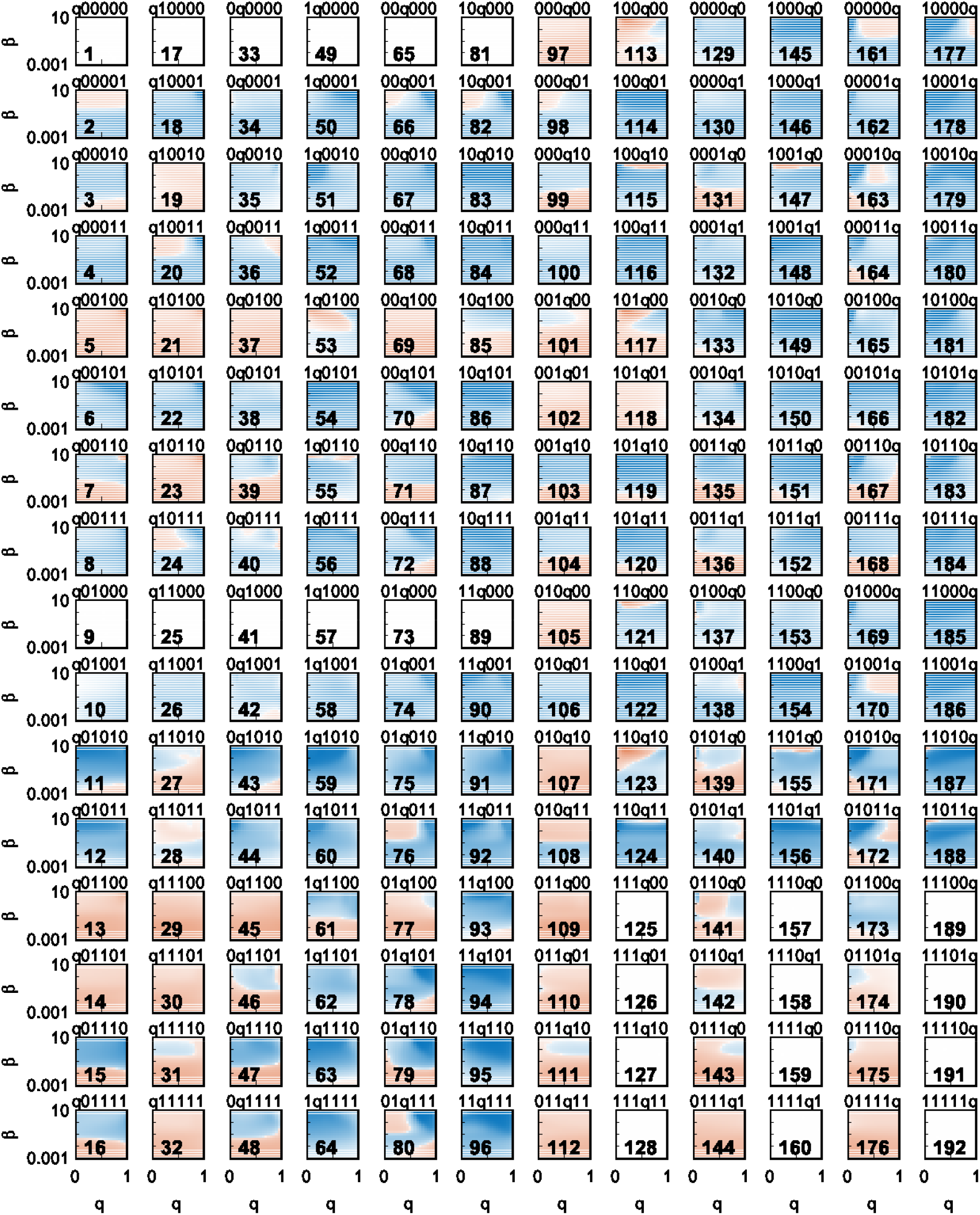
A systematic analysis of the almost-deterministic transition structures for positive selection strength. The figure reproduces the contour plot of Fig.4 (F) for all 192 transition structures in which exactly one transition is probabilistic. There are exactly 24 transition structures for which there is no difference between full and no information for any selection strength. Parameter values: *b*_1_ = 1.8, *b*_2_ = 1.3, *c* = 1, population size *N* = 100, error rate *ε* = 0.01.

**Fig. S9:**
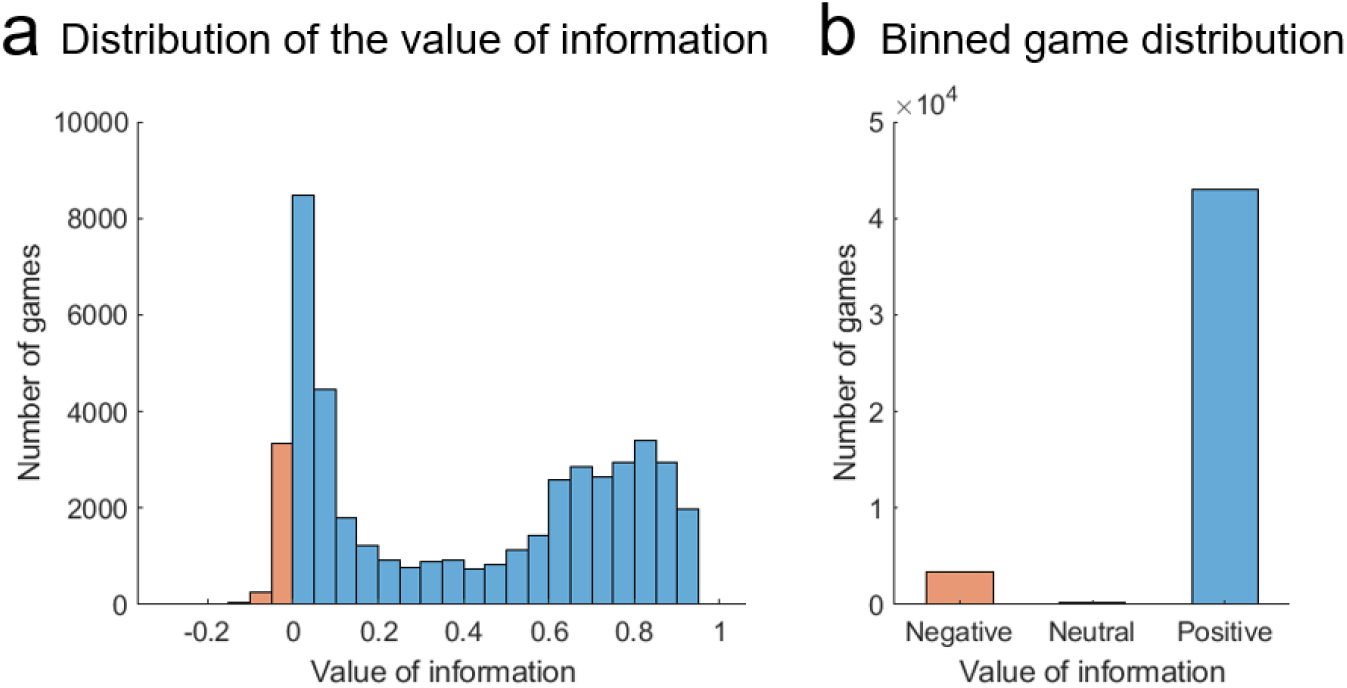
Value of information in games with stochastic transitions. (a) We assume the entries of **q** come from a finite grid 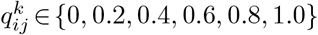 giving rise to 6^6^ = 46, 656 stochastic games. We calculate the average cooperation rates in the games with and without environmental state and compute the value of information. We plot the distribution of the value of information in all stochastic games under consideration. (b) We bin the games in three categories: games with positive (blue), negative (red) and neutral (white) value of information. Parameter values: *b*_1_ = 1.8, *b*_2_ = 1.3, *c* = 1, population size *N* = 100, error rate *ε* = 0.01.

**Fig. S10:**
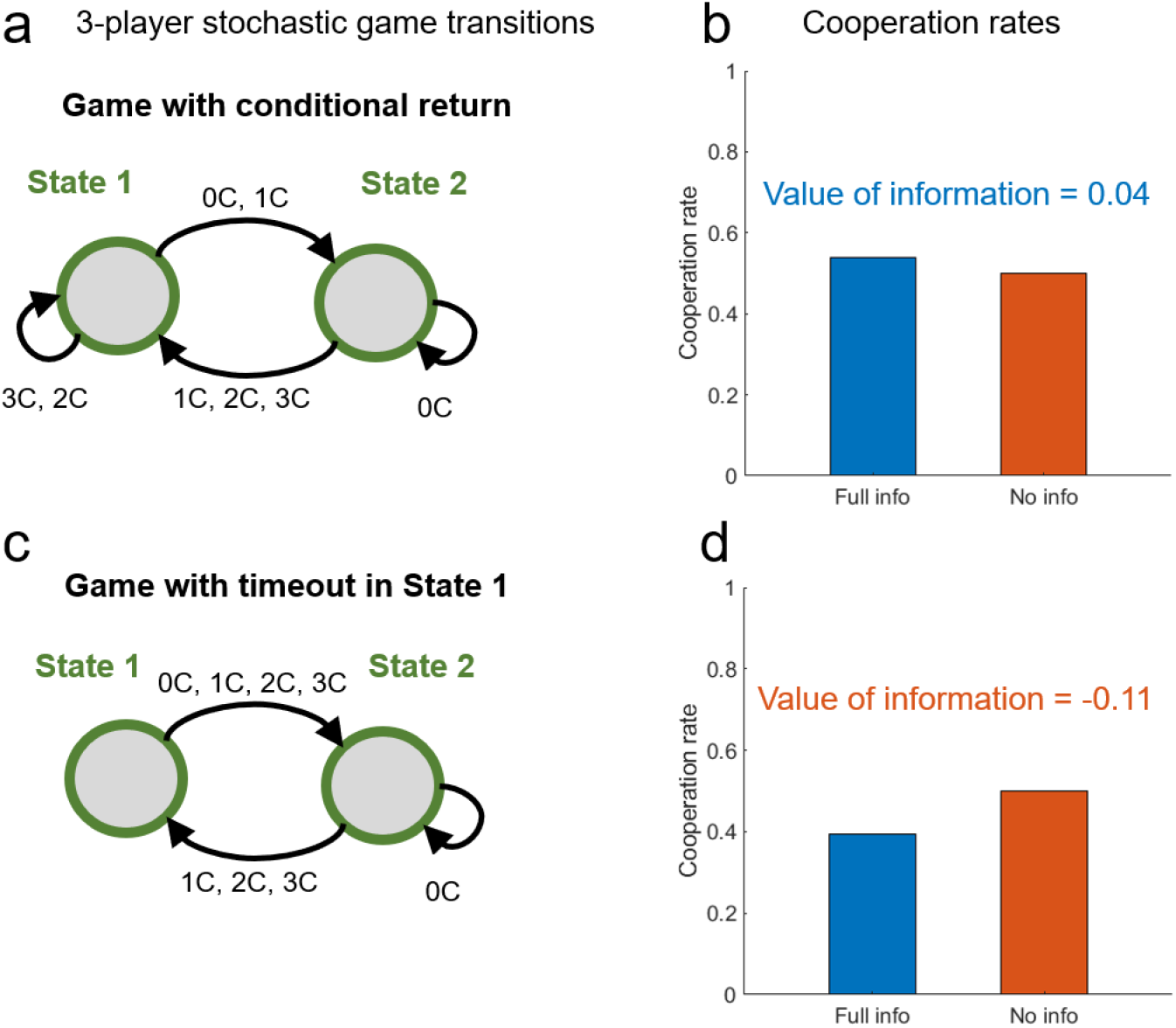
Comparison of the effect of information in two 3-player games. (a-b) We analyse a three-player public good game with a transition vector allowing to return to a better state after defection. We compute cooperation rates in the games with and without environmental information in the limit of no selection, that is, *β* = 0. We find that this transition vector benefits from strategies conditioned on the information about the environmental state. (c-d) We perform the same analysis for a transition vector with a timeout in state 1 and conditional returns to a better state. We find that this game has a benefit of ignorance. Parameter values: *r*_1_ = 1.6, *r*_2_ = 1, *c* = 1, population size *N* = 100, error rate *ε* = 0.01.

## References

[1] Nowak MA. Five rules for the evolution of cooperation. science. 2006;314(5805):1560–1563.

[2] Dugatkin LA. Cooperation among animals: an evolutionary perspective. Oxford Univ. Press; 1997.

[3] Melis AP, Semmann D. How is human cooperation different? Philosophical Transactions of the Royal Society B. 2010;365:2663–2674.

[4] Rand DG, Nowak MA. Human cooperation. Trends in Cogn Sciences. 2012;117:413–425.

[5] Fischbacher U, Gächter S, Fehr E. Are people conditionally cooperative? Evidence from a public goods experiment. Economic Letters. 2001 Apr;71:397–404.

[6] Trivers R. The evolution of reciprocal altruism. The Quarterly Review of Biology. 1971;46(1):35–57.

[7] Hilbe C, Chatterjee K, Nowak MA. Partners and rivals in direct reciprocity. Nature Human Behaviour. 2018;2(7):469–477.

[8] Rapoport A, Chammah AM, Orwant CJ. Prisoner’s dilemma: A study in conflict and cooperation. vol. 165. University of Michigan press; 1965.

[9] Axelrod R. The emergence of cooperation among egoists. American political science review. 1981;75(2):306–318.

[10] Molander P. The optimal level of generosity in a selfish, uncertain environment. Journal of Conflict Resolution. 1985;29:611–618.

[11] Nowak MA, Sigmund K. Tit for tat in heterogeneous populations. Nature. 1992;355(6357):250–253.

[12] Nowak M, Sigmund K. A strategy of win-stay, lose-shift that outperforms tit-for-tat in the Prisoner’s Dilemma game. Nature. 1993;364(6432):56–58.

[13] Kraines DP, Kraines VY. Learning to cooperate with Pavlov an adaptive strategy for the iterated prisoner’s dilemma with noise. Theory and Decision. 1993;35:107–150.

[14] van Segbroeck S, Pacheco JM, Lenaerts T, Santos FC. Emergence of fairness in repeated group interactions. Physical Review Letters. 2012;108:158104.

[15] Pinheiro FL, Vasconcelos VV, Santos FC, Pacheco JM. Evolution of all-or-none strategies in repeated public goods dilemmas. PLoS Comput Biol. 2014;10(11):e1003945.

[16] Hilbe C, Wu B, Traulsen A, Nowak MA. Cooperation and control in multiplayer social dilemmas. Proceedings of the National Academy of Sciences USA. 2014;111:16425–16430.

[17] Stewart AJ, Plotkin JB. From extortion to generosity, evolution in the iterated prisoner’s dilemma. Proceedings of the National Academy of Sciences. 2013;110(38):15348–15353.

[18] Stewart AJ, Plotkin JB. Collapse of cooperation in evolving games. Proceedings of the National Academy of Sciences. 2014;111(49):17558–17563.

[19] Chica M, Hernández JM, Bulchand-Gidumal J. A collective risk dilemma for tourism restrictions under the COVID-19 context. Scientific Reports. 2021;11(1):1–12.

[20] Johnson T, Dawes C, Fowler J, Smirnov O, et al. Slowing COVID-19 transmission as a social dilemma: Lessons for government officials from interdisciplinary research on cooperation. Journal of Behavioral Public Administration. 2020;3(1).

[21] Abel M, Byker T, Carpenter J. Socially optimal mistakes? Debiasing COVID-19 mortality risk perceptions and prosocial behavior. Journal of Economic Behavior & Organization. 2021;183:456–480.

[22] Samuelson CD. Energy conservation: A social dilemma approach. Social Behaviour. 1990;5(4):207–230.

[23] Van Vugt M. Central, individual, or collective control? Social dilemma strategies for natural resource management. American Behavioral Scientist. 2002;45(5):783–800.

[24] Cumming GS. A review of social dilemmas and social-ecological traps in conservation and natural resource management. Conservation Letters. 2018;11(1):e12376.

[25] Vesely S, Klöckner CA, Brick C. Pro-environmental behavior as a signal of cooperativeness: Evidence from a social dilemma experiment. Journal of Environmental Psychology. 2020;67:101362.

[26] Milfont TL. Global warming, climate change and human psychology. Psychological approaches to sustainability: Current trends in theory, research and practice. 2010;19:42.

[27] Tavoni A, Dannenberg A, Kallis G, Löschel A. Inequality, communication, and the avoidance of disastrous climate change in a public goods game. Proceedings of the National Academy of Sciences. 2011;108(29):11825–11829.

[28] Shapley LS. Stochastic games. Proceedings of the National Academy of Sciences. 1953;39(10):1095–1100.

[29] Neyman A, Sorin S, editors. Stochastic games and applications. Dordrecht: Kluwer Academic Press; 2003.

[30] Barfuss W, Donges JF, Kurths J. Deterministic limit of temporal difference reinforcement learning for stochastic games. Physical Review E. 2019;99(4):043305.

[31] Hilbe C, Simsa S, Chatterjee K, Nowak MA. Evolution of cooperation in stochastic games. Nature. 2018;559:246–249.

[32] Su Q, Zhou L, Wang L. Evolutionary multiplayer games on graphs with edge diversity. PLoS computational biology. 2019;15(4):e1006947.

[33] Wang G, Su Q, Wang L. Evolution of state-dependent strategies in stochastic games. Journal of Theoretical Biology. 2021; p. 110818.

[34] Barrett S, Dannenberg A. Sensitivity of collective action to uncertainty about climate tipping points. Nature Climate Change. 2014;4(1):36–39.

[35] Abou Chakra M, Bumann S, Schenk H, Oschlies A, Traulsen A. Immediate action is the best strategy when facing uncertain climate change. Nature communications. 2018;9(1):1–9.

[36] Morton TA, Rabinovich A, Marshall D, Bretschneider P. The future that may (or may not) come: How framing changes responses to uncertainty in climate change communications. Global Environmental Change. 2011;21(1):103–109.

[37] Paarporn K, Eksin C, Weitz JS. Information sharing for a coordination game in fluctuating environments. Journal of theoretical biology. 2018;454:376–385.

[38] Nowak MA, Sasaki A, Taylor C, Fudenberg D. Emergence of cooperation and evolutionary stability in finite populations. Nature. 2004;428(6983):646–650.

[39] Wild G, Traulsen A. The different limits of weak selection and the evolutionary dynamics of finite populations. Journal of Theoretical Biology. 2007;247:382–390.

[40] Wu B, Altrock PM, Wang L, Traulsen A. Universality of weak selection. Physical Review E. 2010;82:046106.

[41] Sigmund K. The calculus of selfishness. vol. 6. Princeton University Press; 2010.

[42] van Veelen M, García J, Rand DG, Nowak MA. Direct reciprocity in structured populations. Proceedings of the National Academy of Sciences USA. 2012;109:9929–9934.

[43] García J, van Veelen M. In and out of equilibrium I: Evolution of strategies in repeated games with discounting. Journal of Economic Theory. 2016;161:161–189.

[44] García J, van Veelen M. No strategy can win in the repeated prisoner’s dilemma: Linking game theory and computer simulations. Frontiers in Robotics and AI. 2018;5:102.

[45] Hilbe C, Martinez-Vaquero LA, Chatterjee K, Nowak MA. Memory-n strategies of direct reciprocity. Proceedings of the National Academy of Sciences USA. 2017;114:4715–4720.

[46] Murase Y, Baek SK. Five rules for friendly rivalry in direct reciprocity. Scientific Reports. 2020;10:16904.

[47] Traulsen A, Pacheco JM, Nowak MA. Pairwise comparison and selection temperature in evolutionary game dynamics. Journal of theoretical biology. 2007;246(3):522–529.

[48] Weitz JS, Eksin C, Paarporn K, Brown SP, Ratcliff WC. An oscillating tragedy of the commons in replicator dynamics with game-environment feedback. Proceedings of the National Academy of Sciences. 2016;113(47):E7518–E7525.

[49] Tilman AR, Plotkin JB, Akçay E. Evolutionary games with environmental feedbacks. Nature Communications. 2020;11(1):1–11.

[50] Wang X, Zheng Z, Fu F. Steering eco-evolutionary game dynamics with manifold control. Proceedings of the Royal Society A. 2020;476:20190643.

[51] Barfuss W, Donges JF, Vasconcelos VV, Kurths J, Levin SA. Caring for the future can turn tragedy into comedy for long-term collective action under risk of collapse. Proceedings of the National Academy of Sciences USA. 2020;117(23):12915–12922.

[52] Hardin G. The Tragedy of the Commons. Science. 1968;162:1243–1248.

[53] Broom M. The Use of Multiplayer Game Theory in the Modeling of Biological Populations. Comments on Theoretical Biology. 2003;8:103–123.

[54] Gokhale CS, Traulsen A. Evolutionary multiplayer games. Dynamic Games and Applications. 2014;4:468–488.

[55] Fudenberg D, Imhof LA. Imitation processes with small mutations. Journal of Economic Theory. 2006;131(1):251–262.

[56] Wu B, Gokhale CS, Wang L, Traulsen A. How small are small mutation rates? Journal of mathematical biology. 2012;64(5):803–827.

[57] Imhof LA, Nowak MA. Stochastic evolutionary dynamics of direct reciprocity. Proceedings of the Royal Society B: Biological Sciences. 2010;277(1680):463–468.

[58] McAvoy A. Comment on “Imitation processes with small mutations”[J. Econ. Theory 131 (2006) 251–262]. Journal of Economic Theory. 2015;159:66–69.

[59] Traulsen A, Hauert C. Stochastic evolutionary game dynamics. Reviews of nonlinear dynamics and complexity. 2009;2:25–61.

