## Supplementary material for "The effect of environmental information on evolution of cooperation in stochastic games": Supplimentary information

Here, we derive results presented in the main manuscript. In **Section 1** we describe the methods used for this research and define the value of information and conditions for when information is valuable and when players can gain the benefit of ignorance. In **Section 2** we provide a systematic analysis of all deterministic stochastic games. First, we provide analytical results for no selection. We provide conditions for when the value of information is positive. Second, we explore how selection strength affects the game dynamics. We then demonstrate how other game parameters affect the value of information. In **Section 3** we extend our systematic analysis to almost-deterministic games and provide additional results on the success of cooperation depending on the information about the environmental state. In **Section 4** we apply our framework to three specific transition vectors to demonstrate how information can alter cooperation rates. In **Mathematical Appendix** we prove all analytical results presented in the main manuscript and Supplementary Information.

### 1 General framework

#### 1.1 Description of the game setup

We explore to which extent conditioning of strategies on the current state of the environment facilitates cooperation in stochastic games. For our baseline model we consider the most simple setup. There are two players who engage in a stochastic game with two possible environmental states. In each state, players can either cooperate (play $C$) or defect (play $D$). Cooperation means to pay a fixed cost $c$ for the co-player to get a benefit $b_i$. This benefit depends on the present state of the environment $s_i \in \{s_1, s_2\}$. Without loss of generality, we assume the first environment is more profitable, such that $b_1 \geq b_2$.

As usual in stochastic games, the next state of the environment may depend on the previous state and on the players' previous actions. Environmental transitions are assumed to be symmetric: they may depend on how many of the players have cooperated in the previous round, but they are independent of who cooperated. We can thus describe the possible transitions by a 6-tuple

$$\mathbf{q} = (q^1_{CC}, q^1_{CD}, q^1_{DD},\ q^2_{CC}, q^2_{CD}, q^2_{DD}). \tag{1}$$

An entry $q^i_{a\tilde{a}}$ gives the probability that the next round's state is $s_1$, given that the previous state was $s_i$ and that the two players chose actions $a$ and $\tilde{a}$, respectively.

We say transitions are *state-independent* if the next state only depends on the players' actions, such that $q^1_{a\tilde{a}} = q^2_{a\tilde{a}}$ for all $a, \tilde{a} \in \{C, D\}$. We call transitions *deterministic* if they are independent of chance events, such that all entries of $\mathbf{q}$ are either zero or one. It follows that there are $2^6 = 64$ deterministic transition vectors. Finally, we call a stochastic game *almost-deterministic* if all entries in $\mathbf{q}$ but one are either zero or one.

Herein, we explore how much cooperation can be sustained when players use memory-1 strategies. Players always know the outcome of the previous round (and are allowed to react to it), but they may or may not account for the environmental state. Specifically, we distinguish two scenarios.

**Full (environmental) information.** In the full-information case, a player's action in the present round may depend on the outcome of the previous round and on the present state of the environment. Such strategies take the form of an 8-tuple,

$$\mathbf{p} = (p^1_{CC}, p^1_{CD}, p^1_{DC}, p^1_{DD},\ p^2_{CC}, p^2_{CD}, p^2_{DC}, p^2_{DD}). \tag{2}$$

An entry $p^i_{a\tilde{a}}$ refers to the player's conditional cooperation probability, depending on the present state $s_i$ and on the player's and co-player's last actions $a$ and $\tilde{a}$, respectively. We refer to the space of all such full-information strategies as $\mathcal{S}_F = [0,1]^8$. For the respective set of pure strategies, we use $\mathcal{P}_F = \{0,1\}^8$.

**No (environmental) information.** In the no-information case, players are still able to condition their behavior on the last round's outcome but not on the present state of the environment. In that case, strategies are given by 4-tuples

$$\mathbf{p} = (p_{CC}, p_{CD}, p_{DC}, p_{DD}). \tag{3}$$

We refer to the space of all such strategies as $\mathcal{S}_N = [0,1]^4$, and to the respective space of pure strategies as $\mathcal{P}_N = \{0,1\}^4$.

We note that no-information strategies can be naturally identified with the subset of full-information strategies for which cooperation probabilities are independent of the present state,

$$\tilde{\mathcal{S}}_F = \left\{ \mathbf{p} \in \mathcal{S}_F \;\middle|\; p^1_{a\tilde{a}} = p^2_{a\tilde{a}} \; \forall \, a, \tilde{a} \in \{C, D\} \right\}. \tag{4}$$

Each strategy in $\mathcal{S}_N$ is mapped to a strategy in $\tilde{\mathcal{S}}_F$ by defining $p^1_{a\tilde{a}} = p^2_{a\tilde{a}} := p_{a\tilde{a}}$.

Herein, we allow for implementation errors. That is, a player who intends to cooperate may defect by mistake with some small probability $\varepsilon$ (analogously, a player who intends to defect my cooperate with the same probability). As a result, a player with strategy $\mathbf{p}$ employs the *effective strategy* $(1-\varepsilon)\mathbf{p} + \varepsilon(\mathbf{1} - \mathbf{p})$. Under this linear transformation, the strategy $ALLD := (0, \ldots, 0)$ becomes $ALLD_\varepsilon = (\varepsilon, \ldots, \varepsilon)$, whereas the strategy $TFT := (1, 0, 1, 0,\ 1, 0, 1, 0)$ becomes $TFT_\varepsilon = (1-\varepsilon, \varepsilon, 1-\varepsilon, \varepsilon,\ 1-\varepsilon, \varepsilon, 1-\varepsilon, \varepsilon)$.

#### 1.2 Calculation of payoffs

In the following, we describe how we can calculate the payoffs when players with arbitrary full-information strategies interact with each other. Since any no-information strategy can be associated with a full-information strategy by Eq. (4), the same method also allows us to compute payoffs when a full-information player is matched with a no-information player, or when two no-information players are matched.

We consider games that are infinitely repeated and in which there is no discounting of the future. Given the players' effective memory-1 strategies $\mathbf{p}$ and $\tilde{\mathbf{p}}$, such games can be described as a Markov chain. The states of this Markov chain correspond to the eight possible outcomes $\omega = (s_i, a, \tilde{a})$ of a given round. Here, $s_i \in \{s_1, s_2\}$ reflects the environmental state, whereas $a, \tilde{a} \in \{C, D\}$ are player 1's and player 2's actions, respectively. The transition probability to move from state $\omega = (s_i, a, \tilde{a})$ in one round to $\omega' = (s'_i, a', \tilde{a}')$ in the next round is given by a product of three factors,

$$m_{\omega,\omega'} = x \cdot y \cdot \tilde{y}. \tag{5}$$

The first factor

$$x = \begin{cases} q^i_{a\tilde{a}} & \text{if } s'_i = s_1 \\ 1 - q^i_{a\tilde{a}} & \text{if } s'_i = s_2 \end{cases} \tag{6}$$

reflects the probability to move from environmental state $s_i$ to $s'_i$, given the player's previous actions. Since the game is symmetric, we note that $q^i_{DC}$ is defined to be equal to $q^i_{CD}$. The other two factors are given by

$$y = \begin{cases} p^{i'}_{a\tilde{a}} & \text{if } a' = C \\ 1 - p^{i'}_{a\tilde{a}} & \text{if } a' = D \end{cases} \quad \text{and} \quad \tilde{y} = \begin{cases} \tilde{p}^{i'}_{\tilde{a}a} & \text{if } \tilde{a}' = C \\ 1 - \tilde{p}^{i'}_{\tilde{a}a} & \text{if } \tilde{a}' = D. \end{cases} \tag{7}$$

They correspond to the conditional probability that each of the two players choses the action prescribed in $\omega'$.

By collecting all these transition probabilities, we obtain an $8 \times 8$ transition matrix $M = (m_{\omega,\omega'})$. Assuming that players are subject to errors and that the game's transition vector satisfies $\mathbf{q} \neq (1, 1, 1, 0, 0, 0)$, this transition matrix has a unique left eigenvector $\mathbf{v}$. The entries $v^i_{a\tilde{a}}$ of this eigenvector give the frequency with which players observe the outcome $\omega = (s_i, a, \tilde{a})$ over the course of the game. For a given transition vector $\mathbf{q}$, we can thus compute the first players' expected payoff as

$$\pi(\mathbf{p}, \tilde{\mathbf{p}}) = b_1\big(v^1_{CC}+v^1_{DC}\big) + b_2\big(v^2_{CC}+v^2_{DC}\big) - c\big(v^1_{CC}+v^1_{CD}+v^2_{CC}+v^2_{CD}\big) \tag{8}$$

The second player's payoff can be computed analogously. We define the average cooperation rate of the two players as

$$\gamma(\mathbf{p}, \tilde{\mathbf{p}}) = \left(v^1_{CC}+\frac{v^1_{CD}+v^1_{DC}}{2}\right) + \left(v^2_{CC}+\frac{v^2_{CD}+v^2_{DC}}{2}\right). \tag{9}$$

In case both players use the same strategy $\mathbf{p}$, we sometimes use the shortcut notation $\gamma(\mathbf{p}) := \gamma(\mathbf{p}, \mathbf{p})$.

#### 1.3 Evolutionary dynamics

To model how players learn to adopt new strategies over time, we study a pairwise comparison process [1] in the limit of rare mutations [2–5].

We consider a population of fixed size $N$. Initially, all players adopt the same resident strategy $\mathbf{p}_R = ALLD$. Then one of the players switches to a randomly chosen alternative strategy $\mathbf{p}_M$. This mutant strategy may then either go extinct or reach fixation eventually, depending on which payoff it yields compared to the resident strategy. If the number of players adopting the mutant strategy is given by $k$, the expected payoffs of the two strategies is

$$\begin{aligned} \pi_R(k) &= \frac{N-k-1}{N-1}\cdot\pi(\mathbf{p}_R, \mathbf{p}_R) + \frac{k}{N-1}\cdot\pi(\mathbf{p}_R, \mathbf{p}_M) \\ \pi_M(k) &= \frac{N-k}{N-1}\cdot\pi(\mathbf{p}_M, \mathbf{p}_R) + \frac{k-1}{N-1}\cdot\pi(\mathbf{p}_M, \mathbf{p}_M) \end{aligned} \tag{10}$$

Based on the players' payoffs, the fixation probability of the mutant strategy can be computed explicitly [6, 7]. It is given by

$$\rho(\mathbf{p}_R, \mathbf{p}_M) = \frac{1}{1+\sum_{i=1}^{N-1}\prod_{k=1}^{i}\exp\Big[-\beta\big(\pi_M(k)-\pi_R(k)\big)\Big]}. \tag{11}$$

Here, $\beta \geq 0$ is the strength of selection. It reflects how important relative payoff advantages are for the evolutionary success of a strategy. If there is no selection and $\beta = 0$, payoffs are completely irrelevant. In that case the fixation probability of a single mutant player in a resident population of size $N$ simplifies to $\rho(\mathbf{p}_R, \mathbf{p}_M) = 1/N$. As $\beta$ increases, the fixation probability is increasingly biased in favor of mutant strategies with a high relative payoff.

If the mutant fixes, it becomes the new resident strategy. Then another mutant strategy is introduced and either fixes or goes extinct. By iterating this basic process for $\tau$ time steps, we obtain a sequence

$(\mathbf{p}_0, \mathbf{p}_1, \mathbf{p}_2, \ldots, \mathbf{p}_\tau)$ where $\mathbf{p}_t$ is the resident strategy present in the population after $t$ mutant strategies have been introduced. Based on this sequence, we can calculate the population's average cooperation rate and payoff as

$$\hat{\pi} = \lim_{\tau\to\infty} \frac{1}{\tau+1} \sum_{t=0}^{\tau} \pi(\mathbf{p}_t, \mathbf{p}_t) \quad \text{and} \quad \hat{\gamma} = \lim_{\tau\to\infty} \frac{1}{\tau+1} \sum_{t=0}^{\tau} \gamma(\mathbf{p}_t, \mathbf{p}_t). \tag{12}$$

In general, these payoff and cooperation averages can only be approximated, by simulating the above described process for a sufficiently long time $\tau$.

Numerically exact results are feasible when mutant strategies are taken from a finite set $\mathcal{P}$. In that case, the evolutionary dynamics can again be described as a Markov chain [2]. Each state of this Markov chain corresponds to one possible resident population $\mathbf{p}\in\mathcal{P}$. Given that the current resident population uses $\mathbf{p}$, the probability that the next resident population uses strategy $\tilde{\mathbf{p}} \neq \mathbf{p}$ is given by $\rho(\mathbf{p}, \tilde{\mathbf{p}})/|\mathcal{P}|$. By calculating the invariant distribution $\mathbf{w} = (w_{\mathbf{p}})$ of this Markov chain, we can compute the average cooperation rates and payoffs according to Eq. (12) by evaluating

$$\hat{\pi} = \sum_{\mathbf{p}\in\mathcal{P}} w_{\mathbf{p}} \cdot \pi(\mathbf{p}, \mathbf{p}) \quad \text{and} \quad \hat{\gamma} = \sum_{\mathbf{p}\in\mathcal{P}} w_{\mathbf{p}} \cdot \gamma(\mathbf{p}, \mathbf{p}). \tag{13}$$

In the following, we report results for the two specific strategy sets $\mathcal{P}_F$ and $\mathcal{P}_N$. By comparing the respective averages $\hat{\pi}_F$, $\hat{\gamma}_F$ with the averages $\hat{\pi}_N, \hat{\gamma}_N$, we wish to characterize for which stochastic games there is an advantage of not knowing the current environmental state.

##### 1.4 The value of information

For a given stochastic game with transition function $\mathbf{q}$ and selection strength $\beta$, we define the value of information $V^\beta(\mathbf{q})$ to be

$$V^\beta(\mathbf{q}) := \gamma_N - \gamma_F. \tag{14}$$

We say there is a *benefit of ignorance* if the value of information is negative, that is, $V^\beta(\mathbf{q}) < 0$. Games with such a benefit of ignorance reflect situations in which players can achieve more cooperation when they are not conditioned on the state they are in.

#### 2 Systematic analysis of all deterministic stochastic games

To make it easier to derive analytical results, we make two further simplifications. (1) We consider *deterministic* transitions that are independent of chance events, such that all entries of $\mathbf{q}$ are either zero or one. There are $2^6 = 64$ deterministic transition vectors. (2) We consider the case of no selection, $\beta = 0$. In the absence of selection, payoffs are irrelevant. This allows us to reduce the number of different cases we need to consider, because we can take the system's symmetries into account.

First, there is no actual difference between “State 1” and “State 2”. So for each stochastic game $\mathbf{q}$, we can define an associated *twin* $\chi(\mathbf{q})$ by relabeling the states,

$$\chi_q(\mathbf{q}) = (1-q^2_{CC}, 1-q^2_{CD}, 1-q^2_{DD},\ 1-q^1_{CC}, 1-q^1_{CD}, 1-q^1_{DD}).$$

Second, for $\beta=0$ it makes no difference to which behavior we refer to as “Cooperation” or “Defection”. It follows that for each stochastic game $\mathbf{q}$ we can define an associated *mirror* game $\psi(\mathbf{q})$ by flipping the meaning of C and D,

$$\psi_q(\mathbf{q}) = (q^1_{DD}, q^1_{CD}, q^1_{CC}, q^2_{DD}, q^2_{CD}, q^2_{CC})$$

Of course, we can also consecutively perform both transformations. This yields a third transformation that we call a mirror-twin,

$$\chi_q \circ \psi_q(\mathbf{q}) = (1-q^2_{DD}, 1-q^2_{CD}, 1-q^2_{CC},\ 1-q^1_{DD}, 1-q^1_{CD}, 1-q^1_{CC}).$$

We can define similar transformation $\chi_p(\mathbf{p})$ and $\psi_p(\mathbf{p})$ for the players’ memory-one strategies, and transformations $\chi_v(\mathbf{v})$ and $\psi_v(\mathbf{v})$ for the resulting invariant distributions. If $\mathbf{v}(\mathbf{p}\,|\,\mathbf{q})$ denotes the invariant distribution of the stochastic game $\mathbf{q}$ between two $\mathbf{p}$-players, we show the following relations (see Lemma 1 and Proposition 1 in **Mathematical Appendix**):

1. $\chi_v\big(\,\mathbf{v}(\chi_p(\mathbf{p})\,|\,\chi_q(\mathbf{q}))\,\big) = \mathbf{v}(\mathbf{p}\,|\,\mathbf{q})$ and $\psi_v\big(\,\mathbf{v}(\psi_p(\mathbf{p})\,|\,\psi_q(\mathbf{q}))\,\big) = \mathbf{v}(\mathbf{p}\,|\,\mathbf{q})$
2. $\gamma\big(\,\mathbf{p}\,|\,\mathbf{q}\,\big) = \gamma\big(\,\chi_p(\mathbf{p})\,|\,\chi_q(\mathbf{q})\,\big)$ and $\gamma\big(\,\mathbf{p}\,|\,\mathbf{q}\,\big) = 1-\gamma\big(\,\psi_p(\mathbf{p})\,|\,\psi_q(\mathbf{q})\,\big)$;
3. Let $\gamma^0_F(\mathbf{q})$ be the average cooperation rate in a full information game with transitions $\mathbf{q}$ and selection strength $\beta = 0$. Then

$$\gamma^0_F(\mathbf{q}) = \gamma^0_F(\chi_q(\mathbf{q})) \quad \text{and} \quad \gamma^0_F(\mathbf{q}) = 1-\gamma^0_F(\psi_q(\mathbf{q})).$$

There are two reasons why these relations are useful. First they imply that with each stochastic game $\mathbf{q}$ that we understand, we immediately understand three other stochastic games, $\chi_q(\mathbf{q})$, $\psi_q(\mathbf{q})$, and $\chi_q \circ \psi_q(\mathbf{q})$. This means that we need to understand fewer distinct cases. Second, in the special case that a transition vector $\mathbf{q}$ is its own mirror, that is $\psi_q(\mathbf{q})=\mathbf{q}$, it follows from the third property above that $\gamma^0_F(\mathbf{q}) = 1/2$. For such transition vectors, information is thus necessarily neutral: the full information game yields the same cooperation rate as the no information game. Similarly, if $\gamma^0_F(\mathbf{q}) < 1/2$ for some transition vector $\mathbf{q}$, then its mirror necessarily has $\gamma^0_F(\psi_q(\mathbf{q})) > 1/2$. That is, for each case with a benefit of ignorance we immediately obtain another case in which there is a disadvantage of ignorance (and vice versa).

In general, we find that among the 64 deterministic games, exactly half of them is neutral. All these cases fall within four possible categories:

(1) The transition vector has an absorbing state: $q^1_{ij}=1$ or $q^2_{ij}=0$ for all $i, j \in \{C, D\}$.

(2) The transition vector is its own mirror: $\psi_q(\mathbf{q}) = \mathbf{q}$.

(3) The transition vector is its own mirror-twin: $\chi_q \circ \psi_q(\mathbf{q}) = \mathbf{q}$.

(4) The transition vector is state-independent: $q^1_{ij} = q^2_{ij}$ for all $i, j \in \{C, D\}$.

We define a simple proxy variable that indicates whether the stochastic game allows players to get absorbed in mutual cooperation or in a mutual defection,

$$X = (q^1_{CC} = 1) + (q^2_{CC} = 0) - (q^1_{DD} = 1) - (q^2_{DD} = 0). \quad (15)$$

In all blue cases in  (cases with no benefit of ignorance) we get $X > 0$, whereas in all red cases (benefit of ignorance), $X < 0$. Moreover, in almost all neutral cases, $X = 0$ (the only exception occurs when the game is neutral because it has an absorbing state). The variable $X$ is computed for all 64 deterministic games in Fig.3b. For increased selection strength in  the cases with the benefit of ignorance are becoming more rare.

### 3 Systematic analysis of all almost-deterministic stochastic games

To explore how pervasive this benefit of ignorance is, we have explored all 192 classes of almost-deterministic games. Fig.S4 illustrates our results for the limit of no selection, whereas Fig.S7 shows the corresponding results for positive selection strength. We call a stochastic game *almost-deterministic* if all entries in $\mathbf{q}$ but one are either zero or one. We first start with analysis of the limit of no selection. Note that there are 24 transition vectors that are absorbing-state vectors for which the benefit of ignorance $V^\beta(q) = 0$ for all $\beta$ and $q$ (see Lemma 2 in **Mathematical Appendix**). Almost all other games follow the same rule as defined in equation (15). However, there is a handful of games that do not follow this rule. We analyse these games in detail here.

#### 3.1 Almost-deterministic cases not explained by $X$

The variable $X$ defined in (15) provides a simple explanation for all deterministic cases. However, it leaves without an explanation behaviour of 24 almost-deterministic cases, for which $X = 0$ and the game does not exhibit neutral behaviour for all $q$. Given the vector transformations we defined earlier, we only need to understand the behaviour of 12 unique cases, where the stochastic transition $q$ occurs only in State 1 of the game. We group these cases based on the similar transitions in State 2 as follows:

(1) $\mathbf{q}_3 = (q00; 010)$ [red] and $\mathbf{q}_{19} = (q10; 010)$ [red];

(2) $\mathbf{q}_6 = (q00; 101)$ [blue] and $\mathbf{q}_{22} = (q10; 101)$ [blue];

(3) $\mathbf{q}_8 = (q00; 111)$ [blue] and $\mathbf{q}_{24} = (q10; 111)$ [blue];

(4) $\mathbf{q}_{10} = (q01; 001)$ [blue], $\mathbf{q}_{26} = (q11; 001)$ [blue] and $\mathbf{q}_{42} = (0q1; 001)$ [yellow];

(5) $\mathbf{q}_{12} = (q01; 011)$ [yellow], $\mathbf{q}_{28} = (q11; 011)$ [red] and $\mathbf{q}_{44} = (0q1; 011)$ [blue].

All of these cases have $X = 0$ but for different values of $q$ some states can become absorbing. For instance, game pairs $\mathbf{q}_6$, $\mathbf{q}_{22}$ and $\mathbf{q}_8$, $\mathbf{q}_{24}$ have an absorbing state $S^1_{CC}$ for $q = 1$, whereas there are no absorbing states in State 2 of the game. Furthermore, neither State 1 nor State 2 of these games possesses a defecting absorbing state. As a result, for $q = 0$ the games are neutral and for $q = 1$ the cooperation rate in the game with full information is higher than in the game with no information.

Note however that it is not a universal rule. For instance, for games $\mathbf{q}_{12}$, $\mathbf{q}_{28}$ and $\mathbf{q}_{44}$ there exists a potentially absorbing state $S^2_{CC}$, which yields a cooperation rate $\gamma = 1$. In order to benefit from it, the game should be able to transit to State 2. For game $\mathbf{q}_{28}$ transition to State 2 is only possible with probability $1 - q$ when both players cooperate in State 1. Otherwise, game 28 can be absorbed in states $S^1_{CC}/S^1_{DD}$ or $S^1_{CD}/S^1_{DC}$, which both yield $\gamma = 1/2$ providing lower cooperation rates. Similar argument applies to game 12 for small $q$. As a result, game 28 and game 12 for small $q$ provide lower cooperation rates than the game with no information. Along the same lines one can analyse the trio $\mathbf{q}_{10}$, $\mathbf{q}_{26}$ and $\mathbf{q}_{42}$, where State 2 of the game provides a chance to be absorbed in the cooperative states $S^2_{CC}$ ($\gamma = 1$) and $S^2_{CD}/S^2_{DC}$ ($\gamma = 1/2$). However, game 42 behaves as games 10 and 26 only for small $q$. For larger $q$, the chances to be absorbed in a defecting state $S^1_{DD}$ are higher, which decreases the level of cooperation (for small $q$ the game is blue and for larger $q$ the game is red).

Perhaps, the most interesting case is the games $\mathbf{q}_3 = (q00; 010)$ and $\mathbf{q}_{19} = (q10; 010)$. Both these games possess $\gamma = 1/2$ for $q = 0$ and $\gamma > 1/2$ for $q = 1$. However, for $q < 1$, the cooperation rates for both these games are $\gamma < 1/2$. For $q$ small, State 1 of the game does not possess any absorbing states and the game always transits to State 2, where it can be absorbed in either $S^2_{CC}/S^2_{DD}$ or $S^2_{CD}/S^2_{DC}$ with $\gamma = 1/2$. For game 3 when $q = 1$, State 1 possesses a potentially absorbing cooperative state $S^1_{CC}$, lifting the total cooperation rate. On the other hand, game 19 for $q = 1$ has a potentially absorbing state $S^1_{CD}/S^1_{DC}$, which lowers its cooperation rate in comparison to game 3.

### 4 Extended analysis transition vectors discussed in the manuscript

#### 4.1 Deterministic game $\mathbf{q}_{39} = (1, 0, 0; 1, 1, 1)$ (timeout game)

The first example we consider is the deterministic game with transition structure $\mathbf{q} = (1, 0, 0; 1, 1, 0)$. In the absence of selection, this game has a cooperation rate of $1/2$ for both games with and without information. However, increasing selection results in higher cooperation rates in the population that conditions their strategies on the environmental information. In order to understand why cooperation rates are different in games with and without information, we explore stability of WSLS strategy.

In the game with full information available to players, for strong selection and parameter values as in Fig.2, we observe that a strategy $\mathbf{p} = (1,0,0,0;x,0,0,1)$ is the most successful (Fig.2c), where $x \in \{0,1\}$. Note that two strategies are possible: either Grim-WSLS given by $\mathbf{p} = (1,0,0,0;1,0,0,1)$ or Grim-risker given by $\mathbf{p} = (1,0,0,0;0,0,0,1)$ analysed in [10]. To explore if this strategy is a subgame perfect Nash equilibrium, we employ the one-shot deviation principle [11]. For this, we calculate continuation payoffs for each of the cases when players either use this strategy or deviate in one round and then return to using this strategy again. Let us first consider payoffs of non-deviating players. Given the nature of memory-1 strategies and state dependency of the game, we need to consider cases of all possible actions in each state.

1. **CC in State 1.** Then, the game transits to State 1 and both players cooperate in all subsequent rounds, that is,

$$\pi_{CC,S1} = b_1 - c.$$

2. **CD/DC in State 1.** Then, the game transits to State 2, where both players will defect. After, the game returns to State 1 and both players still defect as dictates their Grim strategy, which means that after returning to State 2 both players cooperate. After this, the game returns to State 1, where players cooperate in all subsequent rounds. Then, the payoff is given by

$$\pi_{CD,S1} = \delta^2(1-\delta)(b_2 - c) + \delta^3(b_1 - c).$$

3. **DD in State 1.** Here, the game transits to State 2, where both players cooperate and, after returning to State 1, they cooperate in all subsequent rounds, that is,

$$\pi_{DD,S1} = (1-\delta)(b_2 - c) + \delta(b_1 - c).$$

4. **CC in State 2.** This case is similar to CC in State 1 since the game transits to State 1 and both players cooperate in all subsequent rounds, that is,

$$\pi_{CC,S2} = b_1 - c.$$

5. **CD/DC in State 2.** Then, the game transits to State 1, where both players defect. After, the game returns to State 2 and both players cooperate, which recovers cooperation in all subsequent rounds as the game returns to State 1. Then, the payoff is given by

$$\pi_{CD,S2} = \delta(1-\delta)(b_2 - c) + \delta^2(b_1 - c).$$

6. **DD in State 2.** Here, the game transits to State 1, where both players defect, and, via cooperating in State 2, as before, players recover mutual cooperation in all subsequent rounds, that is,

$$\pi_{DD,S2} = \delta(1-\delta)(b_2 - c) + \delta^2(b_1 - c).$$

Now, let us construct the payoffs of the deviating players.

1. **CC in State 1.** In this case a deviating player defects and then returns to using the same strategy as everyone else. That is, the game is in State 1 when the mutant defects and then the game transits to State 2 where both players defect. After the following mutual defection in State 1, players recover cooperation by cooperating in State 2 and cooperate in all subsequent rounds in State 1. That is,

$$\tilde{\pi}_{CC,S1} = (1-\delta)b_1 + \delta^3(1-\delta)(b_2 - c) + \delta^4(b_1 - c).$$

2. **CD/DC in State 1.** Here, the deviation requires the mutant to cooperate in State 2. After transitioning to State 1, both players defect, which again recovers mutual cooperation through transitioning to State 2. The payoff is then given by

$$\tilde{\pi}_{CD,S1} = -(1-\delta)c + \delta^2(1-\delta)(b_2 - c) + \delta^3(b_1 - c).$$

3. **DD in State 1.** As the game transits to State 2, it requires that mutant defects while the second player cooperates. They still recover cooperation after mutual defection in State 1 as before. The payoff of a deviating player is given by

$$\tilde{\pi}_{DD,S1} = (1-\delta)b_2 + \delta^2(1-\delta)(b_2 - c) + \delta^3(b_1 - c).$$

4. **CC in State 2.** This case is similar to CC in State 1 as yields the same payoff to the deviating player

$$\tilde{\pi}_{CC,S2} = (1-\delta)b_1 + \delta^3(1-\delta)(b_2 - c) + \delta^4(b_1 - c).$$

5. **CD/DC in State 2.** The deviating player will cooperate as game transits to State 1 and the second player defects. This leads to the mutual defection in State 2 and a subsequent State 1. After this, both players cooperate in State 2 and all subsequent rounds in State 1. The payoff for this case is given by

$$\tilde{\pi}_{CD,S2} = -(1-\delta)c + \delta^3(1-\delta)(b_2 - c) + \delta^4(b_1 - c).$$

6. **DD in State 2.** Here, the game transits to State 1, where the deviating player is required to cooperate. They again recover mutual cooperation as in the previous case after two rounds of mutual defection. The payoff is then given by

$$\tilde{\pi}_{DD,S2} = -(1-\delta)c + \delta^3(1-\delta)(b_2 - c) + \delta^4(b_1 - c).$$

In order for this strategy to be a subgame perfect Nash equilibrium, we need to check if it always yields higher payoffs than a one-shot deviation. For this, we require that

$$\pi_{CC,S1} \geq \tilde{\pi}_{CC,S1} \quad \text{and} \quad \pi_{CC,S2} \geq \tilde{\pi}_{CC,S2}$$
$$\pi_{CD,S1} \geq \tilde{\pi}_{CD,S1} \quad \text{and} \quad \pi_{CD,S2} \geq \tilde{\pi}_{CD,S2}$$
$$\pi_{DD,S1} \geq \tilde{\pi}_{DD,S1} \quad \text{and} \quad \pi_{DD,S2} \geq \tilde{\pi}_{DD,S2}$$

Note that inequalities for the payoffs after CD in both states are always satisfied. The same is true for the inequality $\pi_{DD,S2} \geq \tilde{\pi}_{DD,S2}$. As payoffs in both states after mutual cooperation are the same, we only need to consider two cases.

1. $\pi_{CC} \geq \tilde{\pi}_{CC}$: this inequality reduces to the following condition

$$(1 + \delta + \delta^2)c \leq \delta(1 + \delta + \delta^2)b_1 - \delta^3 b_2.$$

   For $\delta \to 1$ this inequality is reduced to $3c \leq 3b_1 - b_2$. For the set of parameters we use in Fig. 2, this condition is satisfied.

2. $\pi_{DD,S1} \geq \tilde{\pi}_{DD,S1}$: this inequality can be written as

$$(1 + \delta)c \leq \delta(1 + \delta)b_1 - \delta^2 b_2.$$

   For $\delta \to 1$ it can be reduced to $2c \leq 2b_1 - b_2$. This condition is also satisfied for the set of parameters we chose.

Let us now apply a similar line of argument to strategy WSLS. As can be seen in Fig.2E, this strategy does not evolve in both games with or without information. First, we construct the payoffs for the non-deviating players.

1. **CC in State 1.** All players cooperate in all rounds, that is,

$$\pi_{CC,S1} = b_1 - c.$$

2. **CD/DC in State 1.** Then, players defect in State 2 and recover cooperation in all subsequent rounds in State 1, that is,

$$\pi_{CD,S1} = \delta(b_1 - c).$$

3. **DD in State 1.** The game transits to State 2, where players cooperate, and then they cooperate in all subsequent rounds in State 1,

$$\pi_{DD,S1} = (1 - \delta)(b_2 - c) + \delta(b_1 - c).$$

4. **CC in State 2.** Same as for CC in State 1,

$$\pi_{CC,S2} = b_1 - c.$$

5. **CD/DC in State 2.** Then, players defect in State 1 and recover cooperation in State 2 and cooperate in all subsequent rounds in State 1, that is,

$$\pi_{CD,S2} = \delta(1 - \delta)(b_2 - c) + \delta^2(b_1 - c).$$

6. **DD in State 2.** The game transits to State 1, where players cooperate in all subsequent rounds.

$$\pi_{DD,S2} = b_1 - c.$$

Payoffs of the deviating player then can be calculated in the following way.

1. **CC in State 1.** The deviating player defects in State 1 and after mutual defection in State 2, players recover mutual cooperation in State 1,

$$\tilde{\pi}_{CC,S1} = (1 - \delta)b_1 + \delta^2(b_1 - c).$$

2. **CD/DC in State 1.** The deviating player cooperates in State 2, while the second player defects. This leads to mutual defection in state 1, followed by mutual cooperation in State 2 and subsequent cooperation in all rounds in State 1,

$$\tilde{\pi}_{CD,S1} = -(1 - \delta)c + \delta^2(1 - \delta)(b_2 - c) + \delta^3(b_1 - c).$$

3. **DD in State 1.** The deviating player defects in State 2, which leads to mutual defection in State 1 and recovery of mutual cooperation via State 2,

$$\tilde{\pi}_{DD,S1} = (1-\delta)b_2 + \delta^2(1-\delta)(b_2 - c) + \delta^3(b_1 - c).$$

4. **CC in State 2.** Same as for CC in State 1,

$$\tilde{\pi}_{CC,S2} = (1-\delta)b_1 + \delta^2(b_1 - c).$$

5. **CD/DC in State 2.** The deviating player cooperates in State 1, while the second player defects. This leads to mutual defection in state 2, followed by mutual cooperation in all rounds in State 1,

$$\tilde{\pi}_{CD,S2} = -(1-\delta)c + \delta^2(b_1 - c).$$

6. **DD in State 2.** The deviating player defects in State 1, which leads to mutual defection in State 2 and recovery of mutual cooperation,

$$\tilde{\pi}_{DD,S2} = (1-\delta)b_1 + \delta^2(b_1 - c).$$

As before, we compare all payoffs for deviating and non-deviating players. We see that non-deviating players are better off whenever different actions were played in the previous round independent of the state. Conditions $\pi_{CC} \geq \tilde{\pi}_{CC}$, $\pi_{DD,S1} \geq \tilde{\pi}_{DD,S1}$ and $\pi_{DD,S2} \geq \tilde{\pi}_{DD,S2}$ are reduced to the set of inequalities

$$(1+\delta)c \leq \delta b_1 \quad \text{and} \quad (1+\delta)c \leq \delta(1+\delta)b_1 - \delta^2 b_2$$

While the second condition can re written as $2c \leq 2b_1 - b_2$ for $\delta \to 1$ and is satisfied, the first condition challenges stability of WSLS. As $\delta$ approaches 1, this condition can be written as $2c \leq b_1$. For our set of parameters, this condition is not satisfied, which explains why we rarely observe players adopting this strategy.

By applying a similar argument, we can show that Grim strategy is also an equilibrium. However, the payoff that Grim players obtain in the population of Grim-WSLS or Grim-risker players is lower than the payoff obtained by Grim-WSLS or Grim-risker players among themselves, which explains high frequency of these strategies.

In order to analyse strategies for the game without information, we can consider two cases. First, when players can deduct the current state based on the previous round outcome (assuming they can take into account payoffs they achieve and the actions) but cannot condition their strategy on the current state as such and use this information only for computing the payoffs. This case will be similar in the analysis as the one-shot deviation principle considered for the game with full information. Since WSLS is not an equilibrium strategy, it explains why we mostly observe players using Grim and AllD strategies.

Second, if we assume that players cannot take into account information about previous payoffs and, hence, cannot predict the current state, we analyse the ability of other strategies to invade a population of WSLS players by comparing payoffs mutants can achieve. Note that it is sufficient to consider only strategies like Grim, AllD and risker. The corresponding payoffs $\pi(x, y)$ of a player adopting strategy $x$ against a player adopting strategy $y$ are then given by

$$
\begin{aligned}
\pi(WSLS, WSLS) &= b_1 - c + \mathcal{O}(\epsilon) \\
\pi(Grim, WSLS) &= \frac{1}{35}(15b_1 + 6b_2 - 7c) + \mathcal{O}(\epsilon) \\
\pi(AllD, WSLS) &= \frac{2}{7}b_1 + \frac{3}{14}b_2 + \mathcal{O}(\epsilon) \\
\pi(risker, WSLS) &= \frac{2}{3}b_1 - \frac{1}{3}c + \mathcal{O}(\epsilon)
\end{aligned}
$$

Then, WSLS can withstand an invasion by a population of riskers whenever $2c < b_1$, However, AllD can always invade a population of WSLS players.

##### 4.2 Deterministic game $\mathbf{q}_{38} = (1, 0, 0; 1, 1, 0)$ (timeout game with conditional return)

Deterministic case 38 is characterised by the transition vector $\mathbf{q} = (1, 0, 0; 1, 1, 0)$. Similarly as for the game 39, in the absence of selection, this game has a cooperation rate of $1/2$ for both games with and without information. However, for low to medium selection strength, game with full information demonstrates a higher cooperation rate, while for medium to strong selection, having no information about the current state of the game seems to be more beneficial. In order to understand why the game with full information yields lower cooperation rate, we use similar arguments as we did for the previous game.

There are three most abundant strategies in the game with full information for the parameter set we chose (Fig.2g): $(1, 0, 0, x; y, 0, 0, 1)$, $(1, 0, 1, x; y, 0, 0, 1)$ and $(1, 1, 1, x; y, 0, 1, 0)$, where $x, y \in \{0, 1\}$. All three strategies are self-cooperating and cooperate among each other. However, after consider a one-shot deviation analysis, we see that only the first strategy $(1, 0, 0, x; y, 0, 0, 1)$, which combines WSLS-WSLS, WSLS-risker, Grim-WSLS and Grim-risker strategies, is a subgame perfect Nash equilibrium. One key difference from the game with transition vector $\mathbf{q}_1 = (1, 0, 0;\ 1, 1, 1)$ that makes the WSLS strategy a subgame perfect Nash equilibrium is that mutual defection in State 2 leads to the game remaining in the worse State 2. Then, deviations from the WSLS strategy necessarily yield lower payoff than the payoff of non-deviating players.

Without information, a population consisting of WSLS players cannot be invaded by any other strategy. The payoffs that residents and mutants achieve for the most common non-self-cooperating strategies are given by

$$\pi(WSLS, WSLS) = b_1 - c + \mathcal{O}(\epsilon)$$
$$\pi(Grim, WSLS) = \frac{1}{5}(b_1 + 2b_2 - c) + \mathcal{O}(\epsilon)$$
$$\pi(AllD, WSLS) = \frac{b_2}{2} + \mathcal{O}(\epsilon)$$
$$\pi(risker, WSLS) = \frac{1}{3}(b_1 + b_2 - c) + \mathcal{O}(\epsilon)$$

As can be seen, a population of WSLS players will withstand the invasion of any of these strategies whenever

$$b_2 < 2(b_1 - c).$$

#### 4.3 Almost-deterministic game $\mathbf{q}_{113} = (1, 0, 0; q, 0, 0)$

Next, we consider a stochastic game with transition vector $\mathbf{q} = (1, 0, 0,\ q, 0, 0)$, as displayed in Fig.5. According to this game, players find themselves in the less profitable state 2 if one or both players defected in the previous round. Otherwise, if both players cooperated, they remain in state 1 with certainty if they are already there, or they move towards state 1 with probability $q$ if they start out in state 2.

Numerical simulations depicted in Fig.5 show that there is a benefit of ignorance for a wide range of transition probabilities $q$ and selection strengths $\beta$. We can further support these numerical results by considering the limits of weak and strong selection, respectively.

**Analytical results in the limit of weak selection**

In **Mathematical Appendix**, we derive the following two results:

1. No selection (Propositions 2 and 3 in **Mathematical Appendix**): For $\beta = 0$, no-information yields an average cooperation rate of $\hat{\gamma}_N = 1/2$, whereas full information yields an average cooperation rate of $\hat{\gamma}_F = 1/2 - \frac{3q(1-q)}{64(1+q)}$. In particular, there is a benefit of ignorance $V^0(q) < 0$ for all $q \in (0, 1)$.
2. Weak selection (Propositions 4 and 5 in **Mathematical Appendix**): The above result generalises to positive but sufficiently small selection strengths. That is, there is a threshold $\hat{\beta}$ such that for all $\beta < \hat{\beta}$ there is a benefit of ignorance $V^{\beta}(q) < 0$ for all $q \in (0, 1)$.

**Numerical results in the limit of strong selection**

To get a better insight into the strong selection dynamics, we have characterized all Nash equilibria within the strategy sets $\mathcal{P}_N$ and $\mathcal{P}_F$ as depicted in Fig.5g and Fig.5h.

For no information, we find there are three Nash equilibria for the parameters used in Fig.5. These equilibria are $ALLD = (0,0,0,0)$, $GRIM = (1,0,0,0)$ and $WSLS = (1,0,0,1)$. Out of those, only WSLS is able to sustain cooperation in the presence of rare errors – that is, only for WSLS players we have $\lim_{\varepsilon\to 0}\gamma(\mathbf{p}) = 1$.

For full information, we find 6 distinct equilibria strategies given by the AllD-risker, AllD-Grim, Grim-AllD, Grim-risker, Grim and WSLS strategies. Out of these six equilibria, only WSLS strategy can sustain cooperation in the presence of rare errors.

### Mathematical Appendix

#### Transformations of vectors

**Lemma 1.** *For any memory-one strategy $\mathbf{p} \in S_N$ and transition vector $\mathbf{q}$ we obtain*

$$\gamma(\mathbf{q}|\mathbf{p}) + \gamma(\psi_p(\mathbf{q})|\psi_p(\mathbf{p})) = 1.$$

*Proof.* For every strategy

$$\mathbf{p} = (p^1_{CC}, p^1_{CD}, p^1_{DC}, p^1_{DD}, p^2_{CC}, p^2_{CD}, p^2_{DC}, p^2_{DD})$$

and transition vector

$$\mathbf{q} = (q^1_{CC}, q^1_{CD}, q^1_{DC}, q^1_{DD}, q^2_{CC}, q^2_{CD}, q^2_{DC}, q^2_{DD}) = (\mathbf{q^1}, \mathbf{q}^2)$$

whenever $\mathbf{p}$ plays against itself we shall consider a Markov chain $M(\mathbf{p}, \mathbf{q})$ given by the transition matrix

$$M = \left(\begin{array}{c|c} [\mathbf{q}^1]^T\mathbf{1}\otimes A & [(\mathbf{1}-\mathbf{q}^1)]^T\mathbf{1}\otimes B \\ \hline [\mathbf{q}^2]^T\mathbf{1}\otimes A & [(\mathbf{1}-\mathbf{q}^2)]^T\mathbf{1}\otimes B \end{array}\right),$$

where $\mathbf{q}^T$ is a transpose of the vector $\mathbf{q}$, $\mathbf{1}$ is a vector of ones, $\otimes$ is a Hadamard product and $A$ and $B$ are matrices such that

$$A = \begin{pmatrix} p^1_{CC}p^1_{CC} & p^1_{CC}(1-p^1_{CC}) & (1-p^1_{CC})p^1_{CC} & (1-p^1_{CC})(1-p^1_{CC}) \\ p^1_{CD}p^1_{DC} & p^1_{CD}(1-p^1_{DC}) & (1-p^1_{CD})p^1_{DC} & (1-p^1_{CD})(1-p^1_{DC}) \\ p^1_{DC}p^1_{CD} & p^1_{DC}(1-p^1_{CD}) & (1-p^1_{DC})p^1_{CD} & (1-p^1_{DC})(1-p^1_{CD}) \\ p^1_{DD}p^1_{DD} & p^1_{DD}(1-p^1_{DD}) & (1-p^1_{DD})p^1_{DD} & (1-p^1_{DD})(1-p^1_{DD}) \end{pmatrix}$$

and

$$B = \begin{pmatrix} p^2_{CC}p^2_{CC} & p^2_{CC}(1-p^2_{CC}) & (1-p^2_{CC})p^2_{CC} & (1-p^2_{CC})(1-p^2_{CC}) \\ p^2_{CD}p^2_{DC} & p^2_{CD}(1-p^2_{DC}) & (1-p^2_{CD})p^2_{DC} & (1-p^2_{CD})(1-p^2_{DC}) \\ p^2_{DC}p^2_{CD} & p^2_{DC}(1-p^2_{CD}) & (1-p^2_{DC})p^2_{CD} & (1-p^2_{DC})(1-p^2_{CD}) \\ p^2_{DD}p^2_{DD} & p^2_{DD}(1-p^2_{DD}) & (1-p^2_{DD})p^2_{DD} & (1-p^2_{DD})(1-p^2_{DD}) \end{pmatrix}.$$

Hence, the cooperation rate for $M(\mathbf{p},\mathbf{q})$ is defined as

$$\gamma(\mathbf{q}|\mathbf{p}) = \lim_{\epsilon\to 0}\Big(v^1_{CC} + v^2_{CC} + \frac{v^1_{CD} + v^2_{CD} + v^1_{DC} + v^2_{DC}}{2}\Big),$$

where $\mathbf{v}(\mathbf{p},\mathbf{q})$ is a stationary distribution of $M(\mathbf{p},\mathbf{q})$. The transition matrix for $\phi_{\mathbf{p}}$ is derived as

$$\tilde{M}(\mathbf{p}) = EM(\mathbf{p})E,$$

where $E$ is a permutation matrix given by

$$E = \left(\begin{array}{c|c} \tilde{E} & \mathbf{0} \\ \hline \mathbf{0} & \tilde{E} \end{array}\right),$$

and $\tilde{E}$ is such that

$$\tilde{E} = \begin{pmatrix} 0 & 0 & 0 & 1 \\ 0 & 0 & 1 & 0 \\ 0 & 1 & 0 & 0 \\ 1 & 0 & 0 & 0 \end{pmatrix}.$$

Hence, if $\mathbf{v}(\mathbf{p})$ is a stationary distribution of $M(\mathbf{p})$, then

$$\begin{aligned} &\mathbf{v}(\mathbf{p})M(\mathbf{p}) = \mathbf{v}(\mathbf{p}) \Rightarrow \\ &\mathbf{v}(\mathbf{p})E\tilde{M}(\mathbf{p})E = \mathbf{v}(\mathbf{p}) \Rightarrow \\ &\mathbf{v}(\mathbf{p})E\tilde{M}(\mathbf{p}) = \mathbf{v}(\mathbf{p})E, \end{aligned}$$

and hence $\mathbf{v}(\psi_p(\mathbf{p}) \,|\, \psi_q(\mathbf{q})) := \mathbf{v}(\mathbf{q}|\mathbf{p})E$ is a stationary distribution of $\tilde{M}(\mathbf{p})$. Then, these vectors are defined as

$$\begin{aligned} \mathbf{v}(\mathbf{q}|\mathbf{p}) &= (v^1_{CC}, v^1_{CD}, v^1_{DC}, v^1_{DD}, v^2_{CC}, v^2_{CD}, v^2_{DC}, v^2_{DD}), \\ \mathbf{v}(\psi_p(\mathbf{p}) \,|\, \psi_q(\mathbf{q})) &= (v^1_{DD}, v^1_{DC}, v^1_{CD}, v^1_{CC}, v^2_{DD}, v^2_{DC}, v^2_{CD}, v^2_{CC}). \end{aligned}$$

By the definition of $\gamma(\mathbf{q}|\mathbf{p})$ we obtain

$$\gamma(\mathbf{q}|\mathbf{p}) = \lim_{\epsilon\to 0}\gamma(\mathbf{p}_\epsilon) = \lim_{\epsilon\to 0}\left(v^1_{CC} + v^2_{CC} + \frac{v^1_{CD}+v^1_{DC}+v^2_{CD}+v^2_{DC}}{2}\right),$$
$$\gamma(\psi_p(\mathbf{q})|\psi_p(\mathbf{p})) = \lim_{\epsilon\to 0}\gamma(\psi_p(\mathbf{q})|\psi_p(\mathbf{p}_\epsilon)) = \lim_{\epsilon\to 0}\left(v^1_{DD} + v^2_{DD} + \frac{v^1_{CD}+v^1_{DC}+v^2_{CD}+v^2_{DC}}{2}\right).$$

Hence,

$$\gamma(\mathbf{q}|\mathbf{p}) + \gamma(\psi_p(\mathbf{q})|\psi_p(\mathbf{p})) = \lim_{\epsilon\to 0}\left(\gamma(\mathbf{q}|\mathbf{p}_\epsilon) + \gamma(\psi_p(\mathbf{q})|\psi_p(\mathbf{p}_\epsilon))\right) = \lim_{\epsilon\to 0}\left(\sum_{i=1}^{8} v_i\right) = 1.$$

□

**Proposition 1.** *Consider an arbitrary transition vector* $\mathbf{q}\in[0,1]^6$. *Then for both, full and no information, the following are true*

1. $\hat{\gamma}(\mathbf{q}) = 1 - \hat{\gamma}\big(\psi_q(\mathbf{q})\big)$.
2. $\hat{\gamma}(\mathbf{q}) = \hat{\gamma}\big(\chi_q(\mathbf{q})\big)$.
3. $\hat{\gamma}(\mathbf{q}) = 1 - \hat{\gamma}\big(\psi_q \circ \chi_q(\mathbf{q})\big)$.

*Proof.* **Part 1.** For a transition vector

$$\psi_q(\mathbf{q}) = (q^1_{DD}, q^1_{DC}, q^1_{CD}, q^1_{CC}, q^2_{DD}, q^2_{DC}, q^2_{CD}, q^2_{CC}) = (\psi_q(\mathbf{q^1}), \psi_q(\mathbf{q}^2))$$

we obtain a new Markov chain $M(\mathbf{p}, \psi_q(\mathbf{q}))$ such that

$$M = \left(\begin{array}{c|c} [\psi_q(\mathbf{q}^1)]^T\mathbf{1}\otimes A & [(\mathbf{1}-\psi_q(\mathbf{q}^1))]^T\mathbf{1}\otimes B \\ \hline [\psi_q(\mathbf{q}^2)]^T\mathbf{1}\otimes A & [(\mathbf{1}-\psi_q(\mathbf{q}^2))]^T\mathbf{1}\otimes B \end{array}\right),$$

and the vector $\psi_q(\mathbf{q}^i)$ can be derived as

$$\psi_q(\mathbf{q}^i) = \tilde{E}_1\mathbf{q}^i,$$

where $\tilde{E}_1$ is a permutation matrix such that

$$\tilde{E}_1 = \begin{pmatrix} 0 & 0 & 0 & 1 \\ 0 & 1 & 0 & 0 \\ 0 & 0 & 1 & 0 \\ 1 & 0 & 0 & 0 \end{pmatrix}.$$

Next, we define a global permutation matrix $E_1$ as

$$E_1 = \left(\begin{array}{c|c} \tilde{E}_1 & \mathbf{0} \\ \hline \mathbf{0} & \tilde{E}_1 \end{array}\right).$$

Following a similar argument as in Lemma 1, we obtain that $\mathbf{v}(\mathbf{p}, \mathbf{q})E_1$ is a stationary distribution of $M(\mathbf{p}, \psi_q(\mathbf{q}))$ given by

$$\begin{aligned} \mathbf{v}(\mathbf{p}, \mathbf{q})E_1 &= \left(v^1_{CC}, v^1_{CD}, v^1_{DC}, v^1_{DD}, v^2_{CC}, v^2_{CD}, v^2_{DC}, v^2_{DD}\right) \left(\begin{array}{c|c} \tilde{E}_1 & \mathbf{0} \\ \hline \mathbf{0} & \tilde{E}_1 \end{array}\right) \\ &= \left(v^1_{DD}, v^1_{DC}, v^1_{CC}, v^1_{CC}, v^2_{DD}, v^2_{DC}, v^2_{CD}, v^2_{CC}\right). \end{aligned}$$

Then,

$$\hat{\gamma}(\psi_q(\mathbf{q})) = \lim_{\epsilon \to 0} \Big(v^1_{DD} + v^2_{DD} + \frac{v^1_{DC} + v^2_{DC} + v^1_{CD} + v^2_{CD}}{2}\Big),$$

and hence $\hat{\gamma}(\mathbf{q}) = 1 - \hat{\gamma}(\psi_q(\mathbf{q}))$.

**Part 2.** Part 2 can be shown following a similar logic as in Part 1 using the permutation matrix

$$E_2 = \left(\begin{array}{c|c} \mathbf{0} & \tilde{E}_2 \\ \hline \tilde{E}_2 & \mathbf{0} \end{array}\right),$$

where

$$\tilde{E}_2 = \begin{pmatrix} 1 & 0 & 0 & 0 \\ 0 & 1 & 0 & 0 \\ 0 & 0 & 1 & 0 \\ 0 & 0 & 0 & 1 \end{pmatrix}.$$

Hence,

$$\hat{\gamma}\big(\chi_q(\mathbf{q})\big) = \lim_{\epsilon \to 0} \Big(v^2_{CC} + v^1_{CC} + \frac{v^2_{CD} + v^1_{CD} + v^2_{DC} + v^1_{DC}}{2}\Big),$$

and $\hat{\gamma}(\mathbf{q}) = \hat{\gamma}\big(\chi_q(\mathbf{q})\big)$.

**Part 3.** For Part 3 we use a permutation matrix $E_3$ such that

$$E_3 = E_1 \times E_2 = \left(\begin{array}{c|c} \tilde{E}_1 & \mathbf{0} \\ \hline \mathbf{0} & \tilde{E}_1 \end{array}\right)\left(\begin{array}{c|c} \mathbf{0} & \tilde{E}_2 \\ \hline \tilde{E}_2 & \mathbf{0} \end{array}\right) = \left(\begin{array}{c|c} \mathbf{0} & \tilde{E}_1 \\ \hline \tilde{E}_1 & \mathbf{0} \end{array}\right).$$

Hence,

$$\hat{\gamma}\big(\psi_q \circ \chi_q(\mathbf{q})\big) = \lim_{\epsilon \to 0}\Big(v_{DD}^2 + v_{DD}^1 + \frac{v_{DC}^2 + v_{DC}^1 + v_{CD}^2 + v_{CD}^1}{2}\Big),$$

and $\hat{\gamma}(\mathbf{q}) = 1 - \hat{\gamma}\big(\psi_q \circ \chi_q(\mathbf{q})\big)$.

□

**Lemma 2.** *If the transition vector of the game is such that* $\mathbf{q}^1 = (q_{CC}^1, q_{CD}^1, q_{DD}^1, 0, 0, 0)$ *or* $\mathbf{q}^2 = (1, 1, 1, q_{CC}^2, q_{CD}^2, q_{DD}^2)$*, i.e. either state 1 or 2 is absorbing, then* $V^\beta(q) = 0$ *for all* $\beta$ *and* $q$*.*

*Proof.* If one of the states is absorbing, then players sooner or later will end up in one of the states with the payoff matrix

$$R = \begin{pmatrix} b_i - c & -c \\ b_i & 0 \end{pmatrix}.$$

Then, the Markov chain for such game will be of the form

$$M = \left(\begin{array}{c|c} A & B \\ \hline C & D \end{array}\right),$$

where $C = \mathbf{0}$ for absorbing state 2 and $B = \mathbf{0}$ for absorbing state 1. Hence, for both players, the stationary distribution will be limited to the absorbing state and $v^F = v^N$. Hence, their payoffs and cooperation rates will be the same for any strength of selection and any transition probability $q$.

□

#### Mathematical results for the transition vector $\mathbf{q} = (1, 0, 0; q, 0, 0)$

**Proposition 2.** *When players have no information on the current state, the long-run cooperation rate of the population is* $\bar{\gamma}_N = 1/2$*.*

*Proof.* Due to the bijectivity of $\phi_q(\mathbf{p})$, and Proposition 1, we compute

$$\bar{\gamma}_N = \frac{1}{|S_N|}\sum_{\mathbf{p} \in S_N} \gamma(\mathbf{p}) = \frac{1}{|S_N|}\sum_{\mathbf{p} \in S_N} \frac{\gamma(\mathbf{p}) + \gamma(\phi_q(\mathbf{p}))}{2} = 1/2. \quad (16)$$

□

**Proposition 3.** *When players have full information on the current state, the long-run cooperation rate of the population is*

$$\bar{\gamma}_F = 1/2 - \frac{3q(1-q)}{64(1+q)}.$$

*In particular, $\bar{\gamma}_F \leq 1/2$, with equality if and only if $q = 0$ or $q = 1$. Moreover, the function $\bar{\gamma}_F$ is convex for $0 \leq q \leq 1$ and has a unique minimum at $q^* = \sqrt{2} - 1 \approx 0.41$.*

*Proof.* To get results for full information, we use Mathematica to calculate the self-cooperation rate $\gamma(\mathbf{p})$ for all 32 pure memory-1 strategies in $S_F$ in the limit of rare errors, $\varepsilon \to 0$.

$$\begin{aligned}
&\gamma\big((0,0,0,0,0)\big) = 0, &&\gamma\big((0,1,0,0,0)\big) = 0, &&\gamma\big((1,0,0,0,0)\big) = 0, &&\gamma\big((1,1,0,0,0)\big) = 0,\\
&\gamma\big((0,0,0,0,1)\big) = \tfrac{1}{2}, &&\gamma\big((0,1,0,0,1)\big) = \tfrac{1}{1+q}, &&\gamma\big((1,0,0,0,1)\big) = 1, &&\gamma\big((1,1,0,0,1)\big) = 1,\\
&\gamma\big((0,0,0,1,0)\big) = \tfrac{1}{4}, &&\gamma\big((0,1,0,1,0)\big) = \tfrac{1}{4}, &&\gamma\big((1,0,0,1,0)\big) = \tfrac{1+q}{4}, &&\gamma\big((1,1,0,1,0)\big) = \tfrac{1}{2},\\
&\gamma\big((0,0,0,1,1)\big) = \tfrac{1}{2}, &&\gamma\big((0,1,0,1,1)\big) = \tfrac{3+q}{4+4q}, &&\gamma\big((1,0,0,1,1)\big) = \tfrac{3}{4}, &&\gamma\big((1,1,0,1,1)\big) = \tfrac{3}{4},\\
&\gamma\big((0,0,1,0,0)\big) = \tfrac{1}{4}, &&\gamma\big((0,1,1,0,0)\big) = \tfrac{1}{4}, &&\gamma\big((1,0,1,0,0)\big) = \tfrac{1+q}{4}, &&\gamma\big((1,1,1,0,0)\big) = \tfrac{1}{2},\\
&\gamma\big((0,0,1,0,1)\big) = \tfrac{1}{2}, &&\gamma\big((0,1,1,0,1)\big) = \tfrac{3+q}{4+4q}, &&\gamma\big((1,0,1,0,1)\big) = \tfrac{3}{4}, &&\gamma\big((1,1,1,0,1)\big) = \tfrac{3}{4},\\
&\gamma\big((0,0,1,1,0)\big) = 0, &&\gamma\big((0,1,1,1,0)\big) = 0, &&\gamma\big((1,0,1,1,0)\big) = q, &&\gamma\big((1,1,1,1,0)\big) = 1,\\
&\gamma\big((0,0,1,1,1)\big) = \tfrac{1}{2} &&\gamma\big((0,1,1,1,1)\big) = \tfrac{1}{1+q} &&\gamma\big((1,0,1,1,1)\big) = 1, &&\gamma\big((1,1,1,1,1)\big) = 1.
\end{aligned}$$

By averaging over these 32 self-cooperation probabilities, we obtain the result. □

For the following proofs we require a result from [8]:

**Lemma 3.** *Let $P$ be the stochastic transition matrix with the transition state divided into two sub-states $S = \{S_1, S_2\}$ such that $|S_1| = |S_2|$. This matrix can be written as*

$$P = \left(\begin{array}{c|c} P_{S_1} & P_{S_1S_2} \\ \hline P_{S_2S_1} & P_{S_2} \end{array}\right).$$

*Then, the stationary distribution vector is given by $w = (w_{S_1}, w_{S_2})$ and the following holds*

$$\begin{aligned}
&w_{S_1}\left(P_{S_1} + P_{S_1S_2}(I - P_{S_2})^{-1}P_{S_2S_1}\right) = w_{S_1},\\
&w_{S_2} = w_{S_1}P_{S_1S_2}\left(I - P_{S_2}\right)^{-1}.
\end{aligned}$$

For this section, we will modify notation adopted throughout the manuscript. Let us first construct a Markov chain for the evolutionary dynamics of the game with full information. The state space $S^F$ is given by all possible strategies, that is, vectors $\mathbf{p}^F = (p^1_{CC}, p^2_{CC}, p^2_{CD}, p^2_{DC}, p^2_{DD})$ and $|S| = 32$. Let us order the strategy space in a way that for odd $i$ we obtain $\mathbf{p}^F_{i+1} - \mathbf{p}^F_i = (1,0,0,0,0)$.

Denote this Markov chain as $R^F$ where the off-diagonal elements of the transition matrix are given by

$$\rho_{ij}^F = \frac{1}{1 + \sum_{l=0}^{n-1} \prod_{k=1}^{l} \alpha_k^F}, \tag{17}$$

where $\alpha_k^F = \exp(-\beta(\bar{\pi}_j^F(k) - \bar{\pi}_i^F(k)))$. We define strategy's payoffs, $\pi_i^F$, as

$$\pi_{ij}^F = \sum_{s \in S, a \in A} v_{ij}(s,a) u(s,a),$$

with $v_{ij}(s,a)$ being a Markov chain's stationary distribution and $u(s,a)$ is the corresponding utility function. Then,

$$\bar{\pi}_j^F(k) = \frac{k-1}{n-1}\pi_{jj}^F + \frac{n-k}{n-1}\pi_{ji}^F, \tag{18}$$

$$\bar{\pi}_i^F(k) = \frac{k}{n-1}\pi_{ij}^F + \frac{n-k-1}{n-1}\pi_{ii}^F. \tag{19}$$

**Lemma 4.** *Let the transition probability be $q = 0$. If the two strategies are defined as*

$$\mathbf{p}_i^F = ({}^i p_{CC}^1, {}^i p_{CC}^2, {}^i p_{CD}^2, {}^i p_{DC}^2, {}^i p_{DD}^2)$$

*and*

$$\mathbf{p}_{(i+1)}^F = ({}^{(i+1)} p_{CC}^1, {}^i p_{CC}^2, {}^i p_{CD}^2, {}^i p_{DC}^2, {}^i p_{DD}^2),$$

*then $\pi_{ij}^F = \pi_{(i+1)j}^F, \forall j$, odd $i$, where $\pi_{ij}^F$ is a payoff of strategy $i$ playing against strategy $j$ in the game with full information.*

*Proof.* Consider Markov chains for strategies $i$ and $i+1$, $M_i^F$ and $M_{(i+1)}^F$ respectively. The transition matrices for $q = 0$ are given by

$$M_i^F = \left(\begin{array}{c|c} A_i & B_i \\ \hline 0 & C_i \end{array}\right)$$

and

$$M_{(i+1)}^F = \left(\begin{array}{c|c} A_{(i+1)} & B_i \\ \hline 0 & C_i \end{array}\right).$$

State 2 given by a Matrix $C_i$ is absorbing due to $q = 0$. Then, these Markov chains' stationary distributions, $v_i^F$ and $v_{(i+1)}^F$ are non-zero only on $C_i$. Hence, $v_i^F = v_{(i+1)}^F$. Furthermore, these two games have exactly the same utility functions in the state 2. Hence, $\pi_{ij}^F = \pi_{(i+1)j}^F, \forall j$.

□

**Lemma 5.** *Let the transition probability be $q = 1$. If the two strategies are defined as $\mathbf{p}_i^F = ({}^ip_{CC}^1, {}^ip_{CC}^2, {}^ip_{CD}^2, {}^ip_{DC}^2, {}^ip_{DD}^2)$ and $\mathbf{p}_{(i+1)}^F = ({}^ip_{CC}^1, {}^{(i+1)}p_{CC}^2, {}^ip_{CD}^2, {}^ip_{DC}^2, {}^ip_{DD}^2)$, then $\pi_{ij}^F = \pi_{(i+1)j}^F, \forall j$, odd $i$, where $\pi_{ij}^F$ is a payoff of strategy $i$ playing against strategy $j$ in the game with full information.*

*Proof.* Consider Markov chains for strategies $i$ and $i+1$, $M_i^F$ and $M_{(i+1)}^F$ respectively. The transition matrices for $q = 1$ are given by

$$M_i^F = \left(\begin{array}{c|c} A_i & B_i \\ \hline A_i & B_i \end{array}\right)$$

and

$$M_{(i+1)}^F = \left(\begin{array}{c|c} A_i & B_i \\ \hline A_i & B_i \end{array}\right).$$

Hence, the statement follows.

□

**Proposition 4.** *If a transition probability from state 2 to state 1 is $q = 0$ and $q = 1$, then players gain no benefit from ignorance, that is, $V^\beta(q) = 0$.*

*Proof.* From Lemma 4 for $q = 0$ and Lemma 5 for $q = 1$ we conclude that $\rho_{ij}^F = \rho_{(i+1)j}^F$ and $\rho_{ji}^F = \rho_{j(i+1)}^F, \forall j$ and odd $i$. Note that for cases when $q = 0$ and $q = 1$ strategies are sorted differently, hence, $i$ has different meanings. However, it does not affect the logic of the following proof. Then, the transition matrix of the evolutionary Markov chain can be written as

$$R^F = \begin{pmatrix} \rho_{11}^F & \rho_{11}^F & \rho_{13}^F & \rho_{13}^F & \cdots \\ \rho_{11}^F & \rho_{11}^F & \rho_{13}^F & \rho_{13}^F & \cdots \\ \cdots & \cdots & \cdots & \cdots & \cdots \\ \rho_{31,1}^F & \rho_{31,1}^F & \rho_{31,3}^F & \rho_{31,3}^F & \cdots \\ \rho_{31,1}^F & \rho_{31,1}^F & \rho_{31,3}^F & \rho_{31,3}^F & \cdots \end{pmatrix}.$$

Let us consider the following $32 \times 32$ permutation matrix $E$:

$$E = \begin{pmatrix} 1 & 0 & 0 & 0 & 0 & 0 & 0 & 0 & \dots & 0 & 0 \\ 0 & 0 & 1 & 0 & 0 & 0 & 0 & 0 & \dots & 0 & 0 \\ 0 & 0 & 0 & 0 & 1 & 0 & 0 & 0 & \dots & 0 & 0 \\ 0 & 0 & 0 & 0 & 0 & 0 & 1 & 0 & \dots & 0 & 0 \\ \dots & \dots & \dots & \dots & \dots & \dots & \dots & \dots & \dots & \dots & \dots \\ 0 & 0 & 0 & 0 & 0 & 0 & 0 & 0 & \dots & 1 & 0 \\ 1 & 0 & 0 & 0 & 0 & 0 & 0 & 0 & \dots & 0 & 0 \\ 0 & 0 & 1 & 0 & 0 & 0 & 0 & 0 & \dots & 0 & 0 \\ 0 & 0 & 0 & 0 & 1 & 0 & 0 & 0 & \dots & 0 & 0 \\ 0 & 0 & 0 & 0 & 0 & 0 & 1 & 0 & \dots & 0 & 0 \\ \dots & \dots & \dots & \dots & \dots & \dots & \dots & \dots & \dots & \dots & \dots \\ 0 & 0 & 0 & 0 & 0 & 0 & 0 & 0 & \dots & 1 & 0 \end{pmatrix}.$$

Then,

$$\tilde{R}^F := ER^F E^T = \left(\begin{array}{c|c} A & A \\ \hline A & A \end{array}\right),$$

where $A$ is a $16 \times 16$ matrix with $A_i = (\rho_{(2i-1),1}, \rho_{(2i-1),3}, \dots, \rho_{(2i-1),31})$. The structure of $A$ is such that

$$A = \begin{pmatrix} \rho^F_{11} & \rho^F_{13} & \rho^F_{15} & \cdots & \rho^F_{1,31} \\ \rho^F_{31} & \rho^F_{33} & \rho^F_{35} & \cdots & \rho^F_{3,31} \\ \dots & \dots & \dots & \dots & \dots \\ \rho^F_{29,1} & \rho^F_{29,3} & \rho^F_{29,5} & \cdots & \rho^F_{29,31} \\ \rho^F_{31,1} & \rho^F_{31,3} & \rho^F_{31,5} & \cdots & \rho^F_{31,31} \end{pmatrix}.$$

Assume the Markov chain $\tilde{R}^F$ has a stationary distribution $\tilde{w}^F = (\tilde{w}^F_1, \tilde{w}^F_2)$. Then, by Lemma 3 the following holds

$$\begin{aligned} &\tilde{w}^F_1(A + A(I-A)^{-1}A) = \tilde{w}^F_1, \\ &\tilde{w}^F_2 = \tilde{w}^F_1 A(I-A)^{-1}. \end{aligned}$$

Consider the first equality:

$$\begin{aligned} &\tilde{w}^F_1(A + A(I-A)^{-1}A) = \tilde{w}^F_1 \Rightarrow \\ &\tilde{w}^F_1 A + \tilde{w}^F_1 A(I-A)^{-1}A = \tilde{w}^F_1 \Rightarrow \end{aligned}$$

$$\tilde{w}_1^F A + \tilde{w}_2^F A = \tilde{w}_1^F \Rightarrow$$
$$(\tilde{w}_1^F + \tilde{w}_2^F)A = \tilde{w}_1^F.$$

Next, consider the second equality:

$$\tilde{w}_2^F = \tilde{w}_1^F A(I-A)^{-1} \Rightarrow$$
$$\tilde{w}_2^F(I-A) = \tilde{w}_1^F A \Rightarrow$$
$$\tilde{w}_2^F = \tilde{w}_1^F A + \tilde{w}_2^F A \Rightarrow$$
$$\tilde{w}_2^F = (\tilde{w}_1^F + \tilde{w}_2^F)A.$$

And hence we obtain $\tilde{w}_1^F = \tilde{w}_2^F$ and $2\tilde{w}_1^F$ is a stationary distribution for a Markov chain with the transition matrix $2A$.

Next, we shall consider the game with no information. Let us denote the Markov chain of the evolutionary dynamics in such a game by $R^N$. This game has only 16 strategies due to the fact that $\mathbf{p}^N = (p_{CC}, p_{CD}^2, p_{DC}^2, p_{DD}^2)$. The off-diagonal elements of this matrix are

$$\rho_{ij}^N = \frac{1}{1+\sum_{l=0}^{n-1}\prod_{k=1}^{l}\alpha_k^N},$$

where $\alpha_k^N = \exp(-\beta(\bar{\pi}_i{}^N - \bar{\pi}_j{}^N))$. Let us compare state spaces of Markov chains $R^F$ and $R^N$: the $i$-th state in $R^N$ corresponds to the $(2i-2)$-th state in $R^F$. Given that the utility functions for both games are exactly the same in the state 2 of the stochastic game and that their stochastic games' stationary distributions are equal, we conclude that $\pi_{(2i-1),j}^F = \pi_{ij}^N, \forall j$.

Hence, if the coefficients in the matrices are $\frac{1}{16}$ and $\frac{1}{32}$, then $R^N = 2A$ and $2\tilde{w}_1^F$ is the stationary distribution of $R^N$.

□

**Proposition 5.** *There is a critical selection strength $\beta^* > 0$ such that $V^\beta(q) \geq 0$ for any $\beta < \beta^*$ and any p, with equality if and only if $q = 0$ or $q = 1$.*

*Proof.* We are interested in the value of information denoted as before by

$$V^\beta(q) := \bar{\gamma}_N^\beta(q) - \bar{\gamma}_F^\beta(q),$$

where the average cooperation rates in the full information case and no information case are given by

$$\bar{\gamma}_N^\beta(q) = w_N^\beta \gamma_N(\tilde{\mathbf{p}}),$$
$$\bar{\gamma}_F^\beta(q) = w_F^\beta \gamma_F(\tilde{\mathbf{p}}),$$

with $w_N^\beta$ and $w_F^\beta$ being stationary distributions of the evolutionary processes for the no information and full information cases respectively. Here, $\gamma_N(\tilde{\mathbf{p}}) = (\gamma_N(p_1), \gamma_N(p_2), \dots, \gamma_N(p_{16}))$ and $\gamma_F(\tilde{\mathbf{p}}) = (\gamma_F(p_1), \gamma_F(p_2), \dots, \gamma_F(p_{32}))$ are the cooperation rates for each strategy in the evolutionary strategy space.

Neither $\gamma_N(\tilde{\mathbf{p}})$ nor $\gamma_F(\tilde{\mathbf{p}})$ depend on $\beta$. We need to explore stationary distributions of the evolutionary processes that depend on the payoffs' difference between strategies.

Let us consider a general form of the Markov chain for the game where strategy $\mathbf{p}$ plays against strategy $\mathbf{h}$. Let us denote it as

$$M = \left(\begin{array}{c|c} A & B \\ \hline C & D \end{array}\right),$$

where

$$A = \begin{pmatrix} p^1_{CC}h^1_{CC} & p^1_{CC}(1-h^1_{CC}) & (1-p^1_{CC})h^1_{CC} & (1-p^1_{CC})(1-h^1_{CC}) \\ 0 & 0 & 0 & 0 \\ 0 & 0 & 0 & 0 \\ 0 & 0 & 0 & 0 \end{pmatrix};$$

$$B = \begin{pmatrix} 0 & 0 & 0 & 0 \\ p^2_{CD}h^2_{DC} & p^2_{CD}(1-h^2_{DC}) & (1-p^2_{CD})h^2_{DC} & (1-p^2_{CD})(1-h^2_{DC}) \\ p^2_{DC}h^2_{CD} & p^2_{DC}(1-h^2_{CD}) & (1-p^2_{DC})h^2_{CD} & (1-p^2_{DC})(1-h^2_{CD}) \\ p^2_{DD}h^2_{DD} & p^2_{DD}(1-h^2_{DD}) & (1-p^2_{DD})h^2_{DD} & (1-p^2_{DD})(1-h^2_{DD}) \end{pmatrix};$$

$$C = \begin{pmatrix} qp^1_{CC}h^1_{CC} & qp^1_{CC}(1-h^1_{CC}) & q(1-p^1_{CC})h^1_{CC} & q(1-p^1_{CC})(1-h^1_{CC}) \\ 0 & 0 & 0 & 0 \\ 0 & 0 & 0 & 0 \\ 0 & 0 & 0 & 0 \end{pmatrix};$$

$$D = \begin{pmatrix} (1-q)p^2_{CC}h^2_{CC} & (1-q)p^2_{CC}(1-h^2_{CC}) & (1-q)(1-p^2_{CC})h^2_{CC} & (1-q)(1-p^2_{CC})(1-h^2_{CC}) \\ p^2_{CD}h^2_{DC} & p^2_{CD}(1-h^2_{DC}) & (1-p^2_{CD})h^2_{DC} & (1-p^2_{CD})(1-h^2_{DC}) \\ p^2_{DC}h^2_{CD} & p^2_{DC}(1-h^2_{CD}) & (1-p^2_{DC})h^2_{CD} & (1-p^2_{DC})(1-h^2_{CD}) \\ p^2_{DD}h^2_{DD} & p^2_{DD}(1-h^2_{DD}) & (1-p^2_{DD})h^2_{DD} & (1-p^2_{DD})(1-h^2_{DD}) \end{pmatrix}.$$

For the case with no information, $p^1_{CC} = p^2_{CC}$ and $h^1_{CC} = h^2_{CC}$. Hence, block $B$ is similar for both cases. Let us denote for $\tilde{A}$ the following matrix:

$$\tilde{A} = \begin{pmatrix} p_{CC}^2 h_{CC}^2 & p_{CC}^2(1-h_{CC}^2) & (1-p_{CC}^2)h_{CC}^2 & (1-p_{CC}^2)(1-h_{CC}^2) \\ 0 & 0 & 0 & 0 \\ 0 & 0 & 0 & 0 \\ 0 & 0 & 0 & 0 \end{pmatrix};$$

For the case with no information $A = \tilde{A}$. Further, $C = qA$, $D = B + (1-q)A$ and $\tilde{D} = B + (1-q)\tilde{A}$. Then, the matrices can be rewritten as follows

$$M^N = \left( \begin{array}{c|c} A & B \\ \hline qA & B + (1-q)A \end{array} \right)$$

and

$$M^F = \left( \begin{array}{c|c} A & B \\ \hline qA & B + (1-q)\tilde{A} \end{array} \right).$$

Denote the corresponding stationary distributions by $v^N = (v_1^N, v_2^N)$ and $v^F = (v_1^F, v_2^F)$ separated in blocks in the similar manner as the transition matrices. Then by Lemma 3 the following holds:

$$\begin{cases} v_1^N(A + qB(I - B - (1-q)A)^{-1}A) = v_1^N \\ v_2^N = v_1^N B(I - B - (1-q)A)^{-1}, \end{cases}$$

and

$$\begin{cases} v_1^F(A + qB(I - B - (1-q)\tilde{A})^{-1}A) = v_1^F \\ v_2^F = v_1^F B(I - B - (1-q)\tilde{A})^{-1}. \end{cases}$$

Consider the case when $q = 0$. Then,

$$\begin{cases} v_1^N A = v_1^N \\ v_2^N = v_1^N B(I - B - A)^{-1}, \end{cases}$$

and

$$\begin{cases} v_1^F A = v_1^F \\ v_2^F = v_1^F B(I - B - \tilde{A})^{-1}. \end{cases}$$

However, for the game with no information $A = \tilde{A}$, then we can rewrite conditions for this game as

$$\begin{cases} v_1^N A = v_1^N \\ v_2^N = v_1^N B(I - B - \tilde{A})^{-1}, \end{cases}$$

and hence $v^F = v^N$ and $\pi_{ij}^F = \pi_{ij}^N, \forall i, j$. Hence, for any selection strength $\beta$ and $q = 0$ we obtain $V^\beta(0) = 0$.

Next, let us consider the case when $q = 1$. Then,

$$\begin{cases} v_1^N (A + B(I - B)^{-1} A) = v_1^N \\ v_2^N = v_1^N B(I - B)^{-1}, \end{cases}$$

and

$$\begin{cases} v_1^F (A + B(I - B)^{-1} A) = v_1^F \\ v_2^F = v_1^F B(I - B)^{-1}, \end{cases}$$

and hence $v^F = v^N$ and $\pi_{ij}^F = \pi_{ij}^N, \forall i, j$. Hence, for any selection strength $\beta$ and $q = 1$ we obtain $V^\beta(1) = 0$.

We know that for $q \in (0, 1)$ and $\beta = 0$, $\bar{\gamma}_N(q) = \frac{1}{2}$ and $\bar{\gamma}_F(q) < \frac{1}{2}$. Also, for $\beta = 0$ the evolutionary processes have simple eigenvalues with a corresponding eigenvector $w_F^0$ and $w_N^0$. Hence, by Lemma **??** for $q \in [q_1, q_2]$ there exists $\beta^* > 0$ such that for $\beta < \beta^*$ we obtain $V^\beta(q) > 0$.

On the interval $q \in [0, q_1]$ the slope of the function $\bar{\gamma}_F(q) < 0$ as

$$\frac{\partial}{\partial q} \bar{\gamma}_F(0) \Big|_{\beta=0} = -\frac{3}{64}.$$

Hence, there exists $\beta^* > 0$ such that for $\beta < \beta^*$ we obtain $\frac{\partial}{\partial q} \bar{\gamma}_F(q) \Big|_{\beta} < 0, q \in [0, q_0]$ and hence $V^\beta(q) > 0$.

For the interval $q \in [q_2, 1]$ we obtain $\bar{\gamma}_F(q) > 0$ and, following the similar argument, there exists $\beta^* > 0$ such that for $\beta < \beta^*$ we obtain $B^\beta(q) > 0$.

□

#### References


[1] A. Traulsen, J. M. Pacheco, and M. A. Nowak. Pairwise comparison and selection temperature in evolutionary game dynamics. *Journal of Theoretical Biology*, 246:522–529, 2007.

[2] D. Fudenberg and L. A. Imhof. Imitation processes with small mutations. *Journal of Economic Theory*, 131:251–262, 2006.

[3] L. A. Imhof and M. A. Nowak. Stochastic evolutionary dynamics of direct reciprocity. *Proceedings of the Royal Society B*, 277:463–468, 2010.

[4] B. Wu, C. S. Gokhale, L. Wang, and A. Traulsen. How small are small mutation rates? *Journal of Mathematical Biology*, 64:803–827, 2012.

[5] A. McAvoy. Comment on “imitation processes with small mutations”. *J. Econ. Theory*, 159:66–69, 2015.

[6] M. A. Nowak, A. Sasaki, C. Taylor, and D. Fudenberg. Emergence of cooperation and evolutionary stability in finite populations. *Nature*, 428:646–650, 2004.

[7] A. Traulsen and C. Hauert. Stochastic evolutionary game dynamics. In Heinz Georg Schuster, editor, *Reviews of Nonlinear Dynamics and Complexity*, pages 25–61. Wiley-VCH, Weinheim, 2009.

[8] D. A. Bini, G. Latouche, and B. Meini. *Numerical methods for structured Markov chains*. Oxford University Press on Demand, 2005.

[9] P. D. Lax. *Linear Algebra and Its Applications*. Wiley-Interscience, 2007.

[10] G. Wang, S. Qi, and W. Long *Evolution of state-dependent strategies in stochastic games*. Journal of Theoretical Biology, 527:110818, 2021.

[11] D. Blackwell *Discounted dynamic programming*. The Annals of Mathematical Statistics, 36(1):226–235, 1965.
